# CTCF controls imprinted gene activity at the mouse *Dlk1-Dio3* and *Igf2-H19* domains by modulating allele-specific sub-TAD structure

**DOI:** 10.1101/633065

**Authors:** David Llères, Benoît Moindrot, Rakesh Pathak, Vincent Piras, Mélody Matelot, Benoît Pignard, Alice Marchand, Mallory Poncelet, Aurélien Perrin, Virgile Tellier, Robert Feil, Daan Noordermeer

**Affiliations:** Institute of Molecular Genetics of Montpellier (IGMM), University of Montpellier, CNRS, Montpellier, France; Institute for Integrative Biology of the Cell (I2BC), CEA, CNRS, University Paris-Sud and University Paris-Saclay. Gif-sur-Yvette, France

**Author notes:** Equal author contribution (David Llères, Benoît Moindrot, Rakesh Pathak).

## Abstract

Mammalian genomic imprinting is essential for development and provides a unique paradigm to explore intra-cellular differences in chromatin configuration. Here, we compared chromatin structure of the two conserved imprinted domains controlled by paternal DNA methylation imprints—the *Igf2-H19* and the *Dlk1-Dio3* domains—and assessed the involvement of the insulator protein CTCF. At both domains, CTCF binds the maternal allele of a differentially-methylated region (DMR), in addition to multiple instances of bi-allelic CTCF binding in their surrounding TAD (Topologically Associating Domain). On the paternal chromosome, bi-allelic CTCF binding alone is sufficient to structure a first level of sub-TAD organization. Maternal-specific CTCF binding at the DMRs adds a further layer of sub-TAD organization, which essentially hijacks the existing paternal sub-TAD organisation. Genome-editing experiments at the *Dlk1-Dio3* locus confirm that the maternal sub-TADs are essential during development to maintain the imprinted *Dlk1* gene in an inactive state on the maternal chromosome.

## INTRODUCTION

Genomic imprinting is an epigenetic mechanism through which parental origin dictates mono-allelic expression of about two hundred mammalian genes (1). Imprinting is essential for embryonic development, with diverse disease syndromes in humans attributed to loss of parental specificity (2). The majority of imprinted genes are clustered in about 20 chromosomal domains, where ‘Imprinting Control Regions’ (ICRs) dictate allele-specific gene activity. All ICRs are differentially methylated regions (DMRs) that carry germline-acquired allelic DNA methylation imprints (3).

Most ICRs are methylated on the maternally-inherited chromosome, where they overlap promoters that are expressed only from the paternal chromosome. In contrast, only at two evolutionary conserved domains (*Igf2-H19* and *Dlk1-Dio3*), the ICR is methylated on the paternal allele. Uniquely, these ‘paternal ICRs’ do not overlap promoters but are linked to allele-specific binding of the CTCF insulator protein on the maternal chromosome, either directly to the ICR itself (*Igf2-H19* domain), or to a secondary DMR established early in development (*Dlk1-Dio3* domain) (4–6). Loss of the maternal *Igf2-H19* ICR, or mutations in its CTCF binding sites, lead to the adoption of the paternal transcriptional program, indicating an essential role for allelic CTCF binding (7, 8).

The CTCF insulator protein is essential for the organisation of the genome into ‘Topologically-Associating Domains’ (TADs) (9–11). TADs are 3D structures with enriched intra-domain interactions that tend to insulate genes and their regulatory elements (12). TAD borders are enriched for CTCF binding sites, with a strong enrichment for convergent sites located at both sides of the TAD (9, 13). Disruption of CTCF binding sites at certain, but not all, TAD borders leads to inappropriate activation of surrounding genes during development (14, 15). Within TADs, further levels of chromatin organisation can be observed, sometimes referred to as sub-TADs, with CTCF often being implicated as well (16, 17).

The reported allele-specific binding of CTCF at the paternally imprinted *Igf2-H19* and *Dlk1-Dio3* domains urged us to investigate their chromatin structure within the context of TAD organisation. Previously, non-comprehensive 3C (‘Chromosome Conformation Capture’) studies at the *Igf2-H19* domain reported a number of allele-specific chromatin loops involving the ICR (18, 19). Yet, how these loops are embedded within (sub-)TADs remains unknown due to the incomplete views of DNA contacts and CTCF binding. Moreover, whether the *Dlk1-Dio3* domain adopts a similar allelic 3D architecture, and how chromatin structure is reorganised during imprinted gene activation, remain unexplored. Here, we combined studies of allelic CTCF binding with both high-resolution and single-cell 3D chromatin organization assays to determine the dynamic structuration of the paternally imprinted *Igf2-H19* and *Dlk1-Dio3* domains. Moreover, for the less-characterised *Dlk1-Dio3* domain we performed mechanistic studies to demonstrate the structural and functional importance of allele-specific CTCF binding for correct imprinted gene activation during cellular differentiation.

## RESULTS

### The *Igf2-H19* and *Dlk1-Dio3* domains are located in TADs that include multiple sites of mono- and bi-allelic CTCF binding

To investigate how the *Igf2-H19* and *Dlk1-Dio3* domains are embedded within their respective TADs, we reanalysed high-resolution, but nonallelic, Hi-C data in ESCs. This analysis positioned the *Igf2-H19* and *Dlk1-Dio3* domains within TADs of about 450 kb and 1.6 Mb, respectively [Fig. 1, A and B; (10)]. To address if a parent-of-origin bias may be introduced by allele-specific CTCF binding in these TADs, we performed ChIP-seq on ground-state parthenogenetic (PR8) and androgenetic (AK2) embryonic stem cells (ESCs). For the *Igf2-H19* domain, we detected maternal allele-specific binding of CTCF within the TAD only at the well-characterized ICR located 2-4 kb upstream of the *H19* gene (Fig. 1A, arrow, and fig. S1A). At the *Dlk1-Dio3* domain, our ChIP-seq analysis identified three instances of putative allelic CTCF binding in the TAD on the maternal chromosome (Fig. 1B, arrows). ChIP-seq validation was performed in ESCs (line ‘BJ1’) that were hybrid between the *M. m. domesticus* C57BL/6 and *M. m. molossinus* JF1 inbred lines. This confirmed maternal allele-specific CTCF binding only at the most prominent of these three putative sites (Fig. 1B, solid arrow, and table S1), which we retained for further analysis. This site overlaps the previously identified maternal-specific CTCF binding in humans (5), at a somatic DMR at the first intron of the maternally-expressed *MEG3* lncRNA (*‘Meg3* DMR’; table S2). Closer inspection of this site in the mouse revealed it separated into two peaks that are 900 bp apart (Fig. 1C; sites 1 and 2). ChIP-qPCR experiments in the mono-parental and hybrid ESCs confirmed the robust enrichment of CTCF binding on the maternal chromosome at this site (fig. S1B). Methylation-sensitive endonuclease digestion confirmed methylation of this intronic site on the paternal chromosome, both in hybrid ESCs and in day 9.5 embryos (E9.5) (Fig. 1D and fig. S1, C and D). Both the paternally *H19-Igf2* and *Dlk1-Dio3* imprinted domains are therefore characterized by CTCF recruitment, on the non-methylated maternal allele only, either at the ICR itself [*H19-Igf2* domain; as previously established (4)], or at a close-by secondary DMR (*Dlk1-Dio3* domain; our characterization).

**Figure 1:**
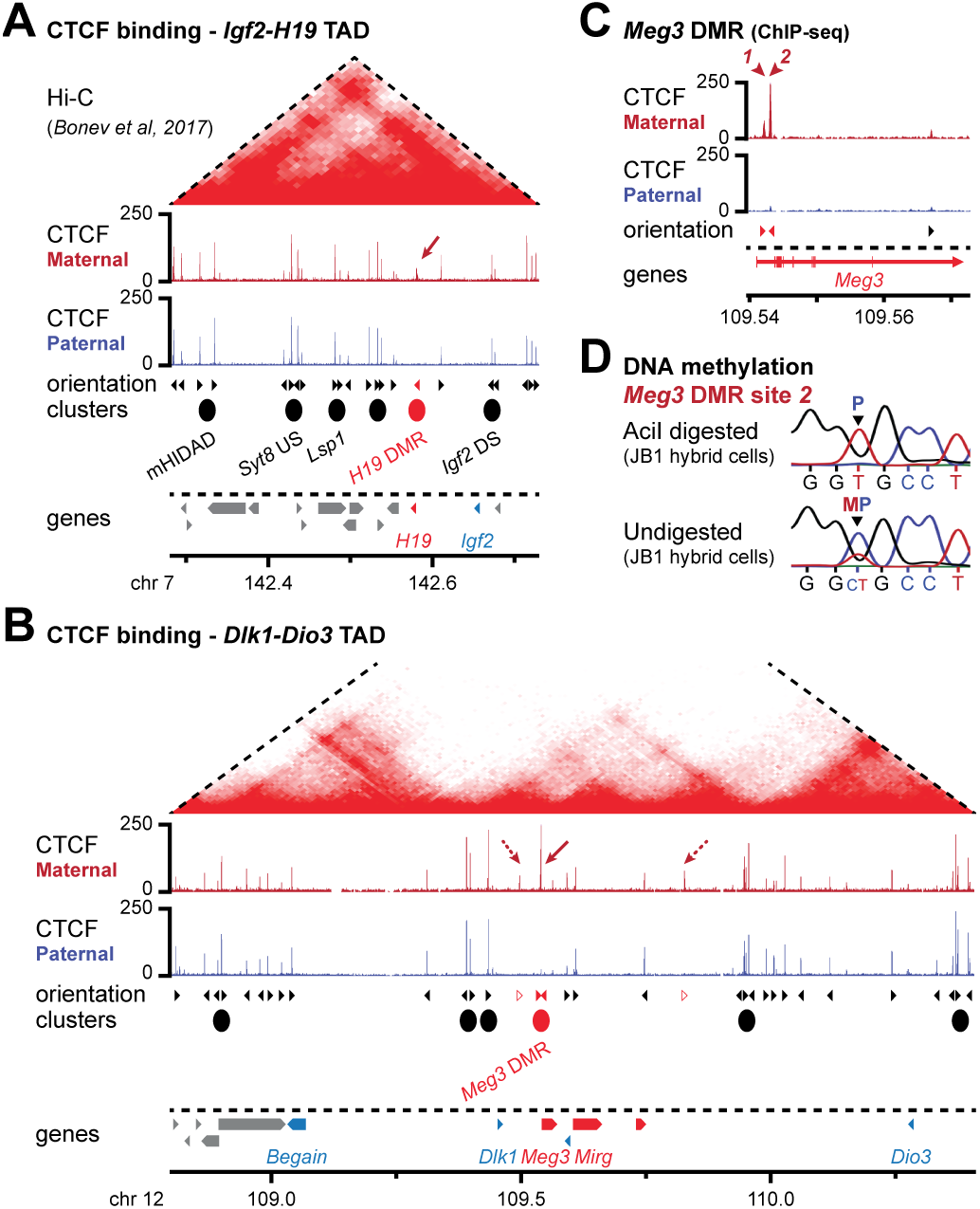
Multiple instances of bi-allelic CTCF binding accompany maternal-specific CTCF binding at the *Meg3* and *H19* DMRs. **A.** CTCF ChIP-seq signal in the TAD containing the *Igf2-H19* domain on the maternal (red) and paternal (blue) chromosome in mono-parental ESCs. The arrow highlights maternal-specific CTCF binding at the *H19* DMR. Reanalysed Hi-C signal is indicated above [(10); 10 kb resolution); the dotted lines indicate TAD boundaries [TADtool; (20)]. The orientation of CTCF sites, the presence of CTCF clusters and genes are indicated below with colours indicating allele-specificity. **B.** CTCF ChIP-seq signal in the TAD containing the *Dlk1-Dio3* domain on the maternal (red) and paternal (blue) chromosome in mono-parental ESCs. The arrow highlights maternal-specific CTCF binding at the *Meg3* DMR. Reanalysed Hi-C signal is indicated above (10); the dotted lines indicate the defined TAD boundaries. **C.** A zoom-in on the *Meg3* DMR CTCF peak reveals it separates into two sub-peaks that are 900 bp apart (sites 1 and 2). **D.** Confirmation of maternal-specific DNA methylation at the *Meg3* DMR in JB1 hybrid ESCs by Sanger sequencing of genomic DNA with (top) and without (bottom) methylation-sensitive *Aci*I digestion. The parental origin of a SNP that distinguishes the maternal and paternal alleles is indicated.

An uncharacterized aspect of both the paternally imprinted domains is extensive additional CTCF binding within their overarching TADs. Our ChIP-seq analysis detected multiple instances of bi-allelic CTCF binding at both the *Igf2-H19* and the *Dlk1-Dio3* domain (Fig. 1, A and B). Within the 450 kb TAD that contains the *Igf2-H19* domain, we particularly noticed the presence of 4 upstream clusters of bi-allelic CTCF binding, positioned between 50 – 250 kb away from the *H19* ICR (Fig. 1A). At the much larger, 1.6 Mb, *Dlk1-Dio3* TAD, we noticed multiple ‘patches’ of bi-allelic CTCF binding as well, including 2 noticeable clusters of CTCF binding around 150 kb upstream of the maternal-specific CTCF-bound DMR (Fig. 1B). These intra-TAD clusters of CTCF binding, together with the maternal CTCF binding at the DMRs, may structure TAD organization at both the domains, or further levels of sub-TAD organization within.

### The *Igf2-H19* and *Dlk1-Dio3* domains are located within non-allele specific TADs

First, we wondered whether the maternal-specific CTCF binding resulted in differences in overall TAD structure at both the paternally imprinted domains. To address this question, we performed high-resolution allele-specific 4C-sequencing using multiple viewpoints in both the *Igf2-H19* and *Dlk1-Dio3 domains* (fig. S2, A and B). As expected, 4C-seq signal obtained for all viewpoints in the domains was largely confined to the TAD that contained the viewpoints themselves, and little signal was detected in neighbouring TADs (fig. S2, C and D). More importantly, a quantitative comparison between the maternal and paternal chromosomes revealed highly similar signal distributions (fig. S2E). We conclude that maternal allele-specific CTCF binding at the *Igf2-H19* and *Dlk1-Dio3* domains does not result in structural changes at the level of overarching TADs.

### Allele-specific sub-TAD organisation of the *Igf2-H19* imprinted domain

To determine if maternal allele-specific CTCF binding may reorganize chromatin organization at the sub-TAD level, we next reassessed our 4C-seq data for the well-characterized *Igf2-H19* domain (fig. S2A). On the maternal chromosome—both in mono-parental and hybrid ESCs—the *H19* DMR strongly interacted with all four upstream clusters of bi-allelic CTCF binding (Fig. 2A, asterisks and fig. S3, A and B). In agreement with the orientation of CTCF binding at the *H19* DMR, all interacting clusters contained at least one binding site orientated towards the *H19* DMR. In contrast, a different configuration was detected at the paternal chromosome, where interactions were globally increased around the viewpoint and towards the 3’ of the *H19* DMR.

**Figure 2:**
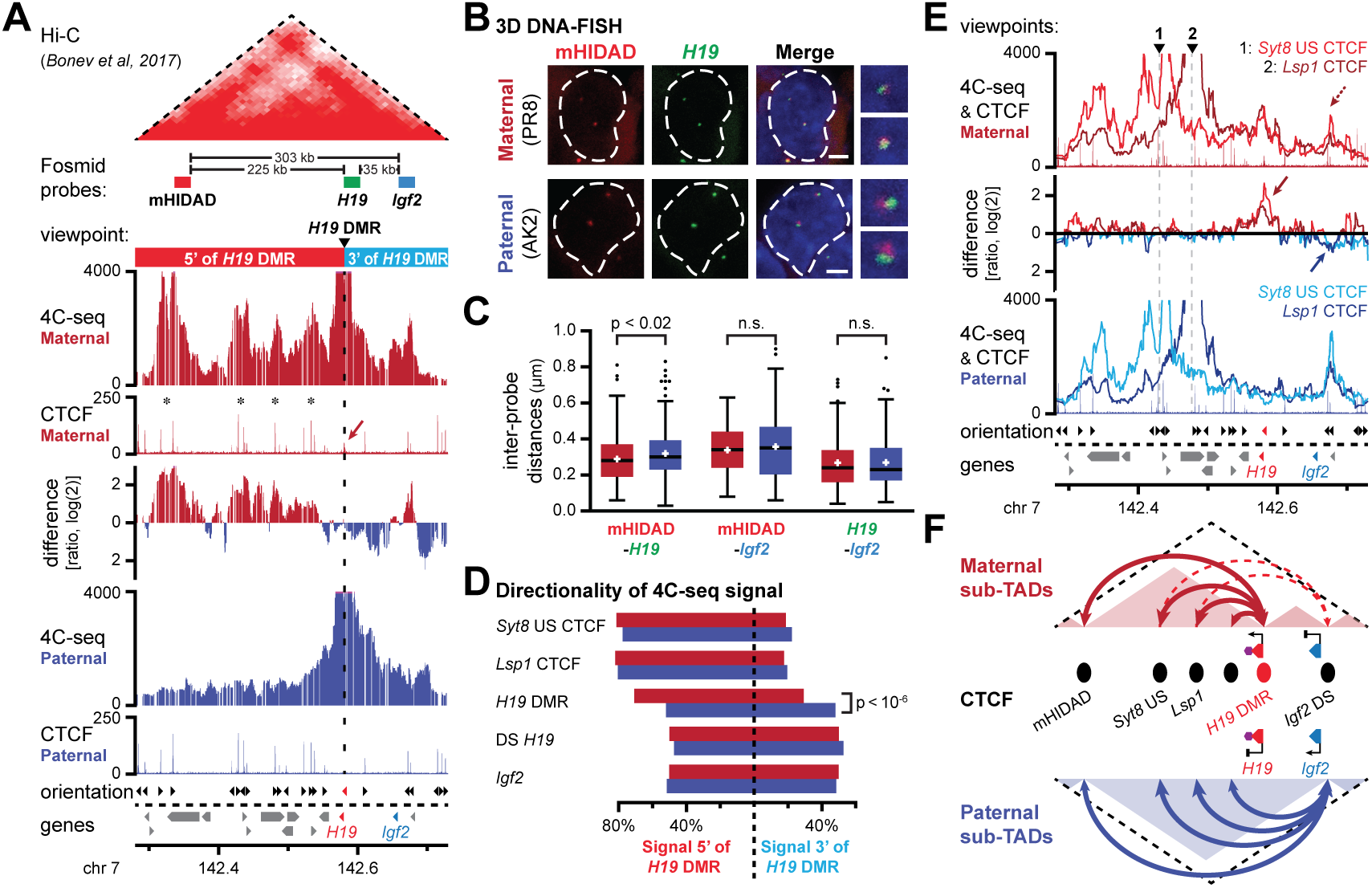
Allele-specific sub-TAD organization of the *Igf2-H19* domain. **A.** 4C-seq signal for the *H19* DMR viewpoint in the *Igf2-H19* TAD on the maternal (red) and paternal (blue) chromosome in mono-parental ESCs. The ratio of maternal/paternal interactions is provided in-between. CTCF ChIP-seq signal is indicated below, with the arrow pinpointing the *H19* DMR. Maternal sub-domains, fosmid probes and the position of the viewpoint are indicated above. **B.** Examples of 3D DNA-FISH with fosmid probes in the *Igf2-H19* TAD (see panel A). Images show representative cells in mono-parental ESCs. Scale bars, 2 μm. **C.** 3D DNA-FISH distance measurements in mono-parental ESCs reveal a smaller distance between the mHIDAD and *H19* probes on the maternal chromosome. **D.** Directionality of 4C-seq signal for indicated viewpoints in the 5’ and 3’ sub-domains (see panel A). **E.** 4C-seq line-graphs for two bi-allelic CTCF clusters (viewpoint 1: upstream *Syt8* cluster and viewpoint 2: *Lsp1* cluster). Arrows indicate increased allele-specific signal at the *H19* DMR (maternal) or downstream of *Igf2* (paternal). **F.** Schematic representation of allele-specific CTCF-structured sub-TAD organization at the *Igf2-H19* domain. CTCF clusters (ovals), allele-specifically expressed genes (triangles) and reported regulatory elements (hexagon) are indicated.

To explore whether the allelic DNA interactions within the *Igf2-H19* TAD correlate with differential higher-order configuration, we measured inter-probe distances by 3D DNA-FISH using fosmid probes (Fig. 2, A and B). Indeed, the average distance between the *H19* and mHIDAD probes was significantly smaller on the maternal chromosome (Fig. 2C and fig. S3C). In contrast, distances between *Igf2* and mHIDAD, a strong bi-allelic CTCF binding site with known structuring function (13), were not significantly different on the maternal and paternal chromosomes. Similarly, we did not detect allelic differences in distance between the *H19* and *Igf2* probes. (Fig. 2C and fig. S3C). Consistently, a 4C-seq viewpoint containing the *Igf2* promoter showed no major allelic differences in its intra-TAD interactions either (fig. S3D). Further reinforcing these observations, statistical analysis of the 4C-seq data confirmed that the parental distribution of signal within the subdomains 5’ and 3’ of the *H19* DMR was significantly different only for the *H19* DMR viewpoint (G-test; Fig. 2D and table S3A). On the maternal chromosome, the CTCF-anchored loops therefore structured a sub-TAD organization that increased the insulation between the reported regulatory elements near the *H19* DMR and the downstream *Igf2* gene (4, 21).

To get a better appreciation of global sub-TAD structure on both the parental chromosomes, we performed further 4C-seq on two of the upstream bi-allelic CTCF clusters (*Syt8* US and *Lsp1* clusters). Whereas both clusters looped towards the *H19* DMR on the maternal chromosome, they formed chromatin loops on the paternal chromosome as well, but now with the more distal bi-allelic CTCF cluster located downstream of the *Igf2* gene (Fig. 2E, arrows and fig. S3E). In the absence of maternal CTCF binding to the *H19* DMR, these upstream CTCF binding clusters thus extended their intra-TAD loops towards further-downstream CTCF sites that are bi-allelic in nature as well. On both the parental chromosomes, the TAD containing the *Igf2-H19* domain therefore has a specific sub-TAD configuration consisting of ensembles of CTCF-anchored chromatin loops. Within this configuration, allele-specific differences are implemented only by CTCF binding at the maternal *H19* DMR (Fig. 2F).

### Recruitment of CTCF to the maternal *Meg3* DMR structures a localised *Dlk1-Meg3* sub-TAD

Next, we shifted our attention to the less characterized *Dlk1-Dio3* domain, which resides within a 1.6 Mb TAD (Fig. 1B). To assess the potential involvement of maternal allele-specific CTCF binding at the *Meg3* DMR, we reassessed our 4C-seq viewpoints targeting both the *Meg3* DMR and the germ-line ICR that acts as a maternal-specific *Meg3* enhancer (the ‘IG-DMR’) (22), as well as the *Dlk1* gene and the CTCF binding site upstream *Dlk1* gene, both in hybrid and mono-parental ESCs (fig. S1A). Consistently, all four viewpoints showed significantly increased interactions on the maternal chromosome, within a 150-kb domain that is demarcated on the right side by the maternal-specific CTCF sites 1 and 2 and on the left by the two bi-allelic clusters of convergently-oriented CTCF clusters (Fig. 3, A and B, and fig. S4A and table S3B). This maternal sub-TAD, which we named the *Dlk1-Meg3* sub-TAD, contains the promoters of *Dlk1* and *Meg3*, both the IG-DMR and the *Meg3* DMR, and several putative *Dlk1* regulatory elements (23). Like for the maternal *Igf2-H19* sub-domain (Fig. 2), the structure of the *Dlk1-Meg3* sub-TAD involves both maternal- and bi-allelic CTCF sites.

**Figure 3:**
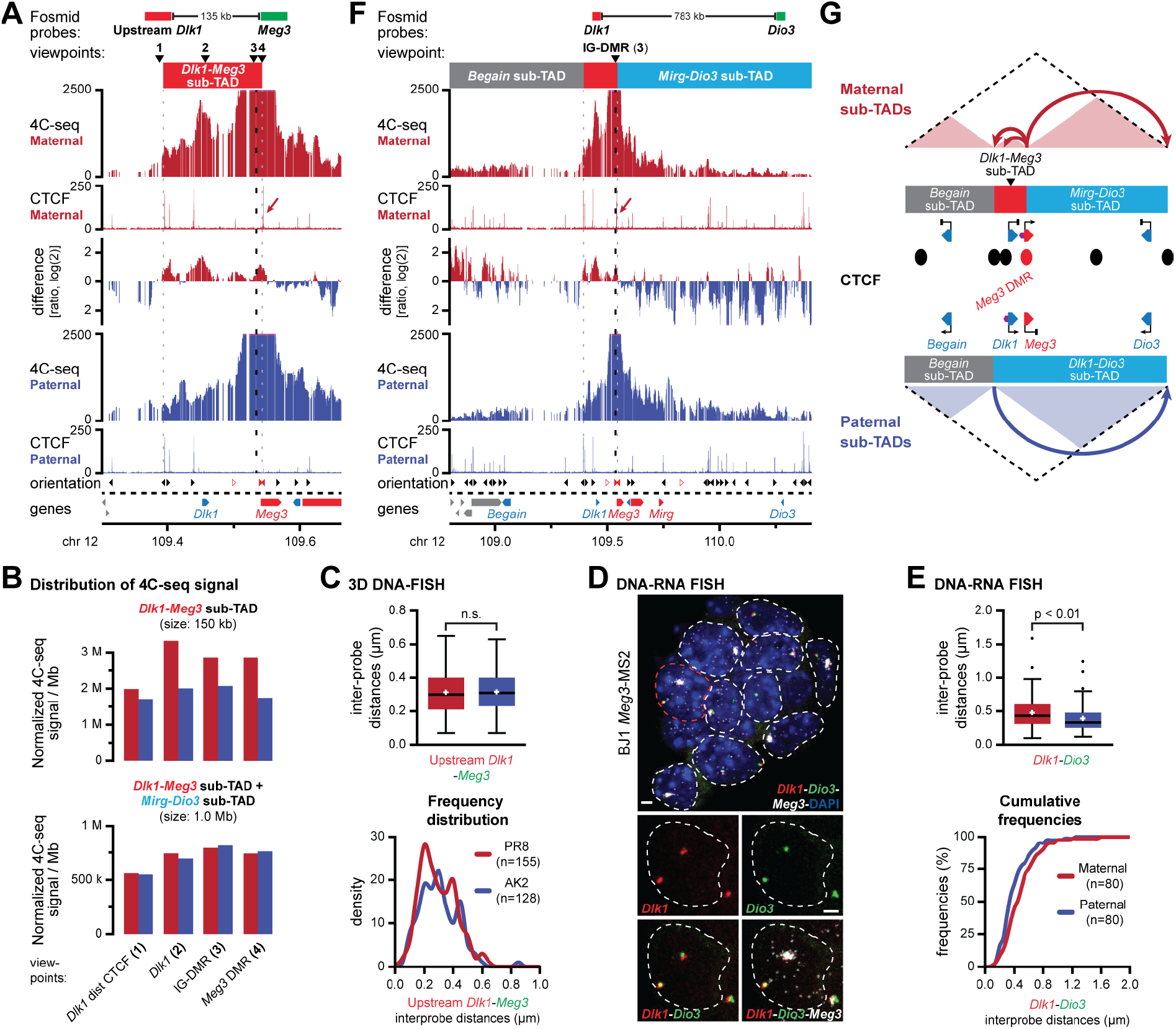
Allele-specific sub-TAD organization of the *Dlk1-Dio3* domain. **A.** 4C-seq signal for the IG-DMR viewpoint in a 300-kb region around *Dlk1-Meg3* on the maternal (red) and paternal (blue) chromosome in hybrid ESCs. The ratio of maternal/paternal interactions is provided in-between. CTCF ChIP-seq signal is indicated below, with the arrow pinpointing the *Meg3* DMR. Fosmid probes, viewpoints and the maternal *Dlk1-Dio3* sub-TAD are indicated above. **B.** Distribution of 4C-seq signal for indicated viewpoints in the *Dlk1-Meg3* sub-TAD (top) and the combined *Dlk1-Meg3* and *Mirg-Dio3* sub-TADs (bottom). **C.** 3D DNA-FISH distance measurements with fosmid probes (see panel **A**) reveal similar distances between both sides of the *Dlk1-Meg3* sub-TAD in mono-parental ESCs. **D.** Allele-specific DNA-RNA FISH with combined fosmids and *Meg3* RNA probes (MS2 sequences) in hybrid ESCs. Images show representative cells. Scale bars, 2 μm. **E.** DNA-RNA FISH distance measurements (see panel **F**) reveal a larger distance between *Dlk1* and *Dio3* on the maternal chromosome. **F.** 4C-seq signal for the IG-DMR viewpoint across the entire *Dlk1-Dio3* TAD. The position of the sub-TADs (red box: *Dlk1-Meg3* sub-TAD; see panel **A**) and fosmid probes are indicated above. **G.** Schematic representation of allele-specific CTCF-structured sub-TAD organization at the *Dlk1-Dio3* domain. CTCF clusters (ovals), allele-specifically expressed genes (triangles) and reported regulatory elements (hexagons) are indicated.

To address how the presence of the *Dlk1-Meg3* sub-TAD correlated with spatial distances between imprinted genes, we performed DNA-FISH using fosmid probes at the borders of this small sub-domain. Inter-probe distances were on average similar between the parental chromosomes (Fig. 3, A and C, and fig. S4B). This finding is similar to our observation that most DNA FISH distances within the imprinted *Igf2-H19* domain are not different between the two parental chromosomes.

Next, we determined how the presence of the *Dlk1-Meg3* sub-TAD on the maternal chromosome influenced contacts between other regions of the imprinted domain. We assayed potential allelic differences in higher-order domain organisation by 3D DNA-FISH in both hybrid and mono-parental ESCs (Fig. 3, D and E, and fig S5A-D). Interestingly, significantly longer distances were measured on the maternal chromosome between all pairs of probes that covered the *Dlk1, Dio3* and *Begain* genes; the three protein-coding genes in the TAD that may acquire mono-allelic expression upon differentiation (24, 25), yet that are inactive in ESCs (table S2). To gain further insight into these differential configurations, we performed 4C-seq in mono-parental ESCs using viewpoints near the *Dlk1, Dio3* and *Begain* promoters (fig. S5E). Replicated experiments revealed no consistent differences between the maternal and paternal chromosomes, including no specific long-range intra-TAD DNA loops on either chromosome beyond the presence of the *Dlk1-Meg3* sub-TAD for the *Dlk1* viewpoint (see also fig. S4A), and a moderate non-allelic enrichment in interactions between the H3K27me3-marked promoters in the domain (26) (fig. S5E, asterisks).

Finally, we determined if the maternal *Dlk1-Meg3* sub-TAD itself engaged in differential contacts with other regions in the TAD. For this purpose, we further analysed our four 4C-seq viewpoints in the *Dlk1-Meg3* sub-TAD (fig. S1A) for differential contacts within the remainder of the TAD. In the upstream subdomain, which we termed the *‘Begain* sub-TAD’, no major difference in total 4C-seq signal between the parental chromosomes was observed. In contrast, in the downstream subdomain, named the *‘Mirg-Dio3* sub-TAD’, 4C-seq signal for all four viewpoints was consistently increased on the paternal allele (Fig. 3F and fig. S4A and S5F). Contacts in the *Mirg-Dio3* sub-TAD therefore displayed an opposite pattern as compared to the maternally-enriched *Dlk1-Meg3* sub-TAD. Whereas this trend was highly significant between the two sub-TADs, their combined signal was essentially the same on the parental chromosomes (Fig. 3B and table S3C). CTCF binding at the maternal chromosome therefore changed the distribution of 4C-seq contacts between the *Dlk1-Meg3* and *Mirg-Dio3* sub-TADs, rather than imposing a complete restructuration of chromatin architecture. This increased insulation may explain the particularly long average DNA-FISH distances on the maternal allele between the *Dio3* probe and probes located in the *Dlk1-Meg3* sub-TAD (Fig. 3E and fig. S5, B and D).

In summary, our combined 4C-seq and DNA-FISH studies in ESCs revealed that the *Dlk1-Dio3* TAD is organised into sub-domains that manifest themselves at multiple levels. The paternal chromosome is structured into the *Begain* sub-TAD and the *Dlk1-Dio3* sub-TAD. On the maternal chromosome, allele-specific CTCF binding at the *Meg3* DMR further divides the *Dlk1-Dio3* sub-domain into the *Dlk1-Meg3* and *Mirg-Dio3* sub-TADs, without affecting the presence of the *Begain* sub-TAD (Fig. 3G). In contrast to the relatively small *Igf2-H19* domain and its host TAD, maternal CTCF binding at the much larger *Dlk1-Dio3* domain and TAD structures a localized sub-domain (*Dlk1-Meg3*), whose insulated nature results in an increased average spatial distance between the *Begain*, *Dlk1* and *Dio3* genes (Fig 3E and fig. S5A-D).

### CTCF binding at the *Meg3* DMR is required for allelic sub-TAD structuration and for correct imprinted activation of *Dlk1*

To confirm if CTCF binding to the maternal *Meg3* DMR is essential for the structure of the maternal *Dlk1-Meg3* sub-TAD and for correct imprinted activation of the *Dlk1* gene, we perturbed CTCF binding using CRISPR-Cas9 genome editing (fig. S6A). We focused on CTCF binding site 2 in the *Meg3* DMR, where CTCF binding is most prominent and conserved in humans (5), and which is oriented towards the CTCF clusters at the 5’ side of the *Dlk1-Meg3* sub-TAD (Fig. 1B). Multiple clonal lines with short bi-allelic deletions comprising site 2 were obtained in both BJ1 and (JF1 × C57BL/6J)F1 mESC (line JB1, fig. S6A). The deletion lines displayed correct imprinted expression of the *Meg3* lncRNA (Fig. 4A and fig. S6B), and unaltered ESC morphology and growth (not shown). Moreover, correct CTCF binding at site 1 and expression of other non-coding RNAs in the domain were maintained in the further characterized hybrid JB1 Δ*site2-cl4* ESCs clone (fig. S6, C and D).

**Figure 4.**
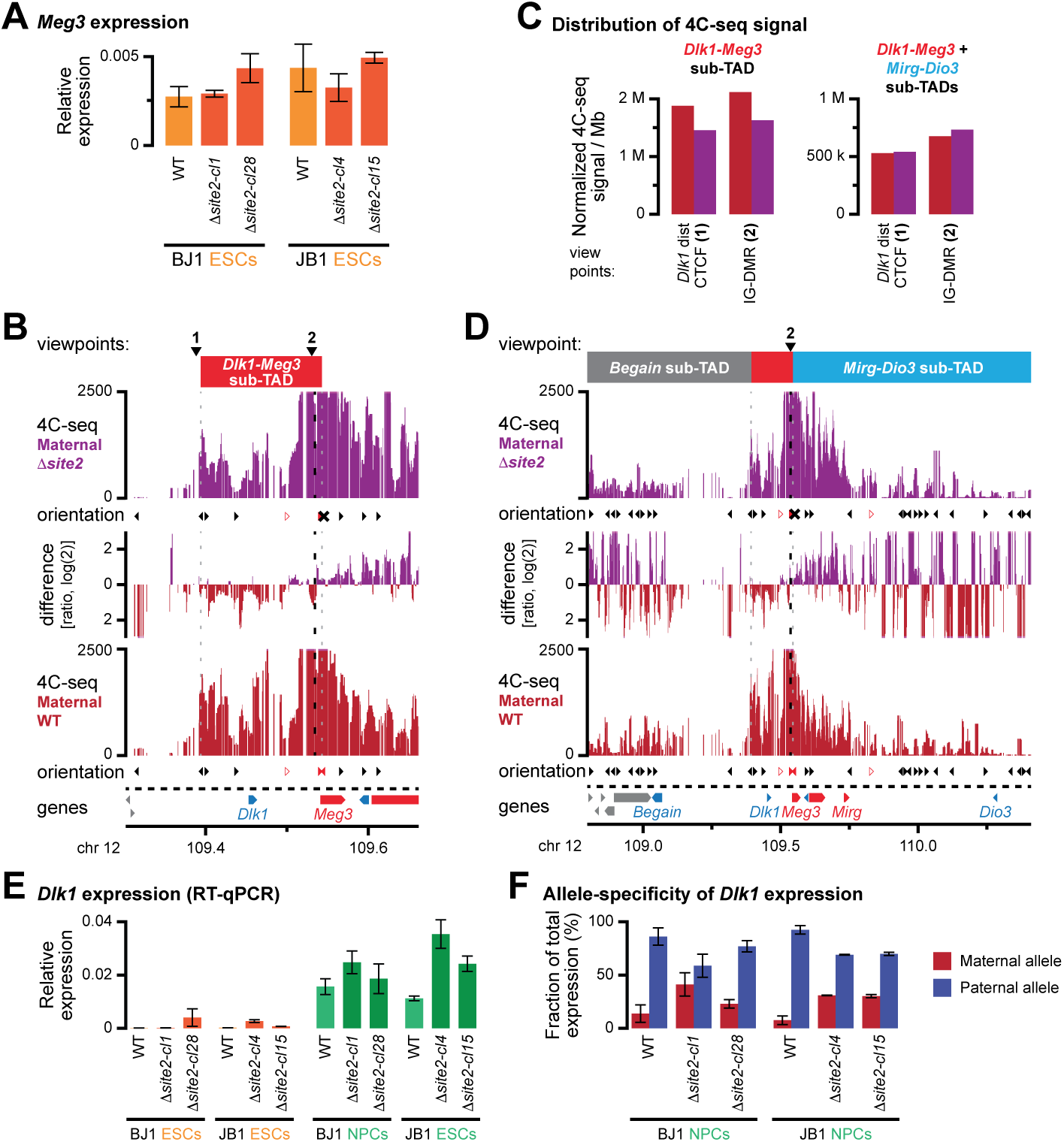
Allelic CTCF binding at the *Meg3* DMR is essential for correct sub-TAD organisation. **A.** *Meg3* expression in hybrid ESCs is maintained upon deletion of CTCF binding site 2 in the *Meg3* DMR, as determined by qRT-PCR. **B.** 4C-seq signal for the IG-DMR (viewpoint **2**) on the maternal alleles in hybrid ESCs with a deleted CTCF site 2 in the *Meg3* DMR (purple) or their WT counterparts (red) in a 300 kb region around the *Dlk1-Dio3* DMRs. The ratio of interactions is provided in-between. The orientation of CTCF sites is indicated below each panel, with the X indicating the deleted CTCF site. Viewpoints and the maternal *Dlk1-Dio3* sub-TAD are indicated above. **C.** Distribution of 4C-seq signal for indicated viewpoints in the *Dlk1-Meg3* sub-TAD (left) and the combined *Dlk1-Meg3* and *Mirg-Dio3* sub-TADs (right). **D.** 4C-seq signal for the IG-DMR viewpoint across the entire *Dlk1-Dio3* TAD. The position of the sub-TADs (red box: *Dlk1-Meg3* sub-TAD) is indicated above. **E.** *Dlk1* expression levels in hybrid ESCs and *in vitro* differentiated NPCs with a deleted CTCF site 2 in the *Meg3* DMR and their WT counterparts, as determined by qRT-PCR. **F.** Allelic *Dlk1* expression becomes relaxed in hybrid NPCs carrying a deletion in CTCF binding site 2 in the *Meg3* DMR, as determined by qRT-PCR.

To determine if allelic CTCF binding directly determined the formation of the *Dlk1-Meg3* sub-TAD, we performed 4C-seq on the JB1 Δ*site2-cl4* ESCs clone. Using viewpoints at the IG-DMR and the CTCF peak upstream of the *Dlk1* gene, we noted that absence of maternal CTCF binding at site 2 strongly reduced interactions within the *Dlk1-Meg3* sub-TAD on the maternal chromosome (Fig. 4, B and C and fig. S6E). Conversely, within the *Mirg-Dio3* sub-TAD, maternal interactions were increased in the mutant cells. Yet, interactions within both sub-TADs combined remained constant (Fig. 4, C and D and fig. S6F). The absence of CTCF binding at *Meg3* DMR site 2 therefore resulted in the adaptation of a more paternal-like 3D architecture on the maternal chromosome.

The domain’s imprinted protein-coding genes, particularly *Dlk1*, are activated upon neural differentiation, but only from the paternal allele (24, 25). To explore if the perturbation of the *Dlk1-Meg3* sub-TAD structure leads to incorrect imprinted activation of the *Dlk1* gene, we performed *in vitro* differentiation of our WT and Δ*site2* hybrid (BJ1 and JB1) ESCs into neural progenitor cells (NPCs) with cortical identity. In this system, imprinted *Dlk1* activation can be readily recapitulated (25, 27). *In vitro* differentiated NPCs from our JB1 and BJ1 deletion ESC lines were developmentally comparable to WT NPCs (fig. S7A). In all lines, *Meg3* remained strictly expressed from the maternal chromosome (fig. S7B). Similar to WT cells, *Dlk1* expression strongly increased upon neural differentiation of our Δ*site2* hybrid cells (Fig. 4E). Deletion of CTCF binding at the *Meg3* DMR site 2 therefore did not interfere with differentiation into NPCs, with persistence of maternal *Meg3* expression or with *Dlk1* activation. In contrast, in all deletion lines, we observed transcriptional activation not only from the paternal chromosome, but also from the maternal chromosome. Allele-specific qRT-PCR revealed that on average 31% of total *Dlk1* transcripts in our deletion NPC-lines were of maternal origin (Fig. 4F and fig. S7B). This partial relaxation of imprinted *Dlk1* activation establishes that CTCF binding to site 2 in the *Meg3* DMR, which instructs the formation of the *Dlk1-Meg3* sub-TAD, prevents activation of the *Dlk1* gene from the maternal chromosome during neural differentiation.

### *Dlk1* activation upon neural differentiation occurs without major restructuring of sub-TAD organisation

To explore whether the allelic, CTCF-mediated sub-TAD organisation contributes directly to the imprinted activation of *Dlk1*, we assessed whether it is maintained in the *in vitro* differentiated NPCs. Re-analysis of published non-allelic Hi-C data, using the same *in-vitro* differentiation protocol of ESCs into NPCs, revealed a similar chromatin architecture of the *Dlk1-Dio3* TAD, with only—as previously reported—a minor increase in long-range intra-TAD interactions upon *in vitro* differentiation (10) (Fig. 5A, green shading, and fig. S8A). We performed allelic 4C-seq on ESC-derived hybrid NPCs using viewpoints at the IG-DMR and the bi-allelic CTCF peak upstream of the *Dlk1* gene (Fig. 5B and fig. S8, B and C). Similar to ESCs, we observed allele-specific differences in the distribution of 4C-seq signal between the *Dlk1-Meg3* and *Mirg-Dio3* sub-TADs, with the total amount of signal in the combined sub-TADs being similar between the parental chromosomes (Fig. 5C, fig. S8D and table S3D). *In vitro* differentiation was therefore not accompanied by drastic reorganisation of sub-TAD configuration. Direct comparison between NPCs and ESCs, on either the paternal or maternal chromosome, revealed no major domain-wide changes in chromatin contacts either (Fig. 5B and fig. S8C). Moreover, comparison of 3D distances between *Dlk1* and *Dio3* using DNA-FISH, although nonallele specific, revealed no significant change in the relative distance between sub-TADs upon differentiation either (fig. S7E). Any changes detected in the 4C-seq analysis therefore do not represent a major reorganization of sub-TAD configuration between NPCs and ESCs. Rather, the *Dlk1-Dio3* sub-TAD organisation remains largely stable during differentiation, and may thus provide allele-specific scaffolding that is required for the correct imprinted activation of *Dlk1* during stem cell differentiation.

**Figure 5:**
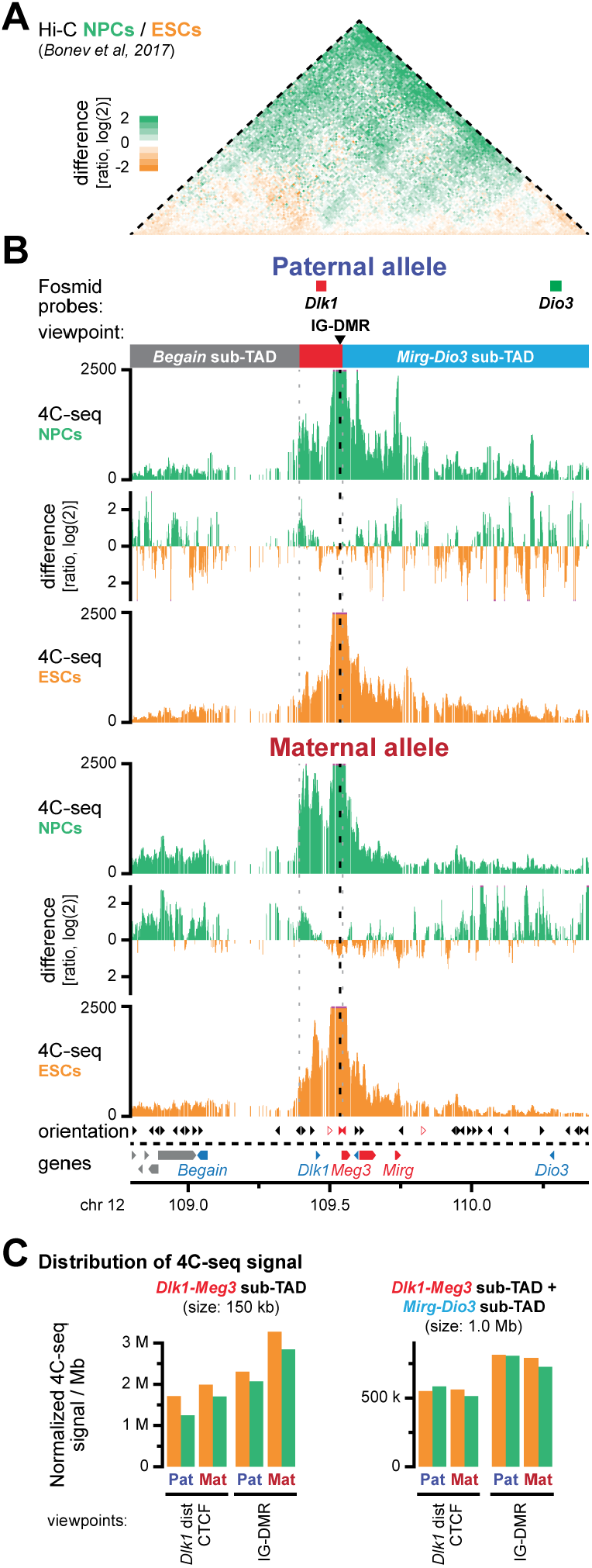
Paternal *Dlk1* activation during neural differentiation occurs without major intra-TAD reorganisation. **A.** Differential non-allelic Hi-C signal upon differentiation of ESCs (orange) to NPCs (green). Hi-C data from reference (10). **B.** 4C-seq signal for the IG-DMR viewpoint in *in vitro* differentiated hybrid NPCs and ESCs on the paternal (top) and maternal (bottom) chromosome in the *Dlk1-Dio3* TAD. The positions of fosmid probes, the viewpoint and sub-TADs (red box: *Dlk1-Meg3* sub-TAD) are indicated above. **C.** Distribution of 4C-seq signal for indicated viewpoints in the *Dlk1-Meg3* sub-TAD (left) and the combined *Dlk1-Meg3* and *Mirg-Dio3* sub-TADs (right).

## DISCUSSION

In this study, we dissected chromatin structure of the two conserved paternally imprinted domains—*Igf2-H19* and *Dlk1-Dio3*—and their surrounding TADs, using genomics and imaging-based approaches. Both domains show maternal-specific CTCF binding at DMRs, together with multiple sites of bi-allelic CTCF binding, to structure localized maternal allele-specific sub-TADs. Unlike the allele-specific chromosome organisation in X-chromosome inactivation (13, 28), these sub-structures are contained within overarching TADs that are similar on both the parental chromosomes. Rather, at both imprinted domains, maternal allele-specific CTCF binding hijacks an existing TAD organisation that is formed between bi-allelic CTCF bound clusters. The resulting maternal chromosome-specific sub-TADs are already established in ESCs, before imprinted activation of protein-coding genes on the paternal alleles (25). These allele-specific sub-TADs may thus provide the ‘instructive’ or ‘permissive’ context for correct developmentally regulated imprinted gene expression during development (29).

More generally, our study highlights striking mechanistic similarities between the two paternally imprinted domains conserved in mammals. At both the domains, sperm-derived DNA methylation imprints mediate, directly or indirectly, maternal allele-specific binding of CTCF to key regions. This structures sub-TADs on the maternal chromosome, which prevents the developmental activation of essential protein-coding genes (*Igf2* at *H19-Igf2*, and *Dlk1* at *Dlk1-Dio3*) on the maternal chromosome. Perturbation of maternal CTCF binding at the DMRs results in loss-of-imprinting of the paternally expressed protein-coding genes, as determined in this study (*Meg3* DMR) or previously published (*H19* DMR) (7). Although the maternal sub-TADs at both paternally imprinted domains are directly associated with lncRNA expression, these sub-TADs may thus ultimately have evolved to repress the developmentally regulated activation of the paternally expressed genes by overriding the action of the inherent paternal 3D organisation. As such, this allele-specific sub-TAD architecture, within a context of overarching non-allelic TADs (30), importantly advances the interpretation of non-comprehensive 3C and CTCF-binding data previously obtained at the *Igf2-H19* domain (18, 19).

How the observed sub-TAD structuration achieves gene repression *in cis*, is different between the two domains though. At the *Igf2-H19* domain, the maternal sub-TADs increase the insulation between the *Igf2* gene and regulatory elements that can activate both *Igf2* and *H19* (21) (Fig. 2F). This mechanism is further supported by a previous study, where positioning of the enhancers downstream of the maternal sub-TADs resulted in the inversion of imprinted gene activation (31). As such, these sub-TADs function as ‘instructive’ chromosomal neighbourhoods that delineate gene-enhancer contacts (14, 29). Yet, their presence or absence is tuned through an epigenetic switch within the context of normal developmental and parental origin.

In contrast, at the *Dlk1-Dio3* domain, the maternal *Dlk1-Meg3* sub-TAD further clusters the repressed *Dlk1* gene and its identified regulatory elements (Fig. 3G). The necessity for this maternal-specific clustering, mediated by CTCF binding to the *Meg3* DMR, was shown by deletion of CTCF binding site 2, which resulted in a more relaxed imprinting of the *Dlk1* gene, with developmental activation now occurring on both parental chromosomes. Whether presence of the maternal *Dlk1-Meg3* sub-TAD similarly represses the imprinted activation of the *Begain* and *Dio3* genes remains to be determined. Both genes are located at considerable distance within the large TAD, and within different sub-TADs (Fig. 3G). These genes do not become activated upon *in vitro* neuronal differentiation, and are not imprinted in *in vitro* generated cortical NPCs, complicating the study of their imprinted activation (25, 27). Recently, we found that the maternal expression of *Meg3*, and possibly the lncRNA itself, prevents activation of *Dlk1* as well (25). We hypothesise that the two mechanisms are linked, with either the *Dlk1-Meg3* sub-TAD focusing or constraining the repressive function of the *Meg3* expression or transcript, or conversely, with maternal *Meg3* expression facilitating CTCF recruitment or its stability of binding at the DMR, possibly through direct RNA-CTCF contacts (32, 33).

Our high-resolution 4C-seq and DNA-FISH studies measure different aspects of chromatin organisation (34, 35). Joint consideration of both types of data can therefore provide complementary, and sometimes apparently paradoxical, insights into the mechanistic and functional organization of chromatin domains. At the *Dlk1-Dio3* domain, our 4C-seq studies identified the presence of the *Dlk1-Meg3* sub-TAD, yet our DNA-FISH studies revealed larger distances on the maternal chromosome between *Dlk1* and *Dio3* (Fig. 3 and fig. S5). Rather than a less compacted maternal *Dlk1-Dio3* TAD, an intra-TAD 3D architecture may be formed where the *Dlk1-Meg3* sub-TAD loops away from the other sub-domains. This is supported by our 3-way DNA-FISH studies where distances involving the central *Dlk1-Meg3* sub-TAD are longer than those between the flanking *Begain* and *Mirg-Dio3* sub-domains (fig. S5D).

At the *Igf2-H19* domain, our DNA-FISH studies revealed that average distances between loci on the parental chromosomes only differ for the *H19* DMR and an upstream CTCF cluster. This appears at odds with our 4C-seq data for the same viewpoint, that shows a highly different pattern of chromatin loops uniquely formed on the maternal chromosome. Interestingly though, for viewpoints at the *H19* DMR and the 5’-located CTCF sites we noticed a considerable enrichment of 4C-seq signal at the *Igf2* gene on the maternal allele as well (Fig 2E, dotted arrow). The *H19* DMR therefore appears unable to impose the observed sub-TAD organisation in all cells (or at all times), possibly due to its relatively low level of CTCF binding (Fig. 1A). This in turn may explain the reported incomplete maternal repression of *Igf2* (36). We speculate that the array of 4 CTCF binding sites at the maternal *H19* DMR (4) hijacks 3D organization at the domain by acting as a ratchet for loops with the upstream bi-allelic CTCF sites, similarly as reported for the *Dxz4* region on the inactive X-chromosome (37), but that this is insufficient to fully override the inherent paternal organisation structured by the loops between the bi-allelic CTCF sites at *Igf2* and the same upstream CTCF sites. At the *Dlk1-Dio3* domain, a similar mechanism may structure the maternal *Dlk1-Meg3* sub-TAD. Here, the CTCF-bound maternal *Meg3* DMR interacts with two separate clusters of bi-allelic CTCF sites in the proximal part of the domain.

In conclusion, our study reveals the importance of maternal-specific CTCF binding to structure a further layer of sub-TAD organization that is essential to override the inherent paternal organization and associated gene activation. Similarly as for the *H19* DMR—where epigenetic alterations that affect CTCF binding cause the growth-related imprinting disorders Beckwith-Wiedemann Syndrome (BWS) and Silver-Russell Syndrome (SRS) (2)—maternal CTCF binding at the *Meg3* DMR is evolutionarily conserved in humans (5). Microdeletions and gains of methylation within this region have recently been linked to the developmental imprinting disorder Kagami-Ogata Syndrome (KOS14) (2, 38), indicating that our observations are relevant to humans as well.

## ACKNOWLEDGMENTS

We thank Patricia Cavelier and Claire Dupont for ESC culture and karyotyping, Yan Jaszczyszyn and the I2BC Next Generation Sequencing Facility, Mylène Docquier and the University of Geneva iGE3 Genomics Platform and Michael Girardot for generating sequencing data, the ‘Montpellier Ressources Imagerie’ platform for advice on microscopy and FACS, and the Feil and Noordermeer labs for useful discussion. 4C-seq data analysis was performed at the Vital-IT Center for high-performance computing (www.vital-it.ch) of the Swiss Institute of Bioinformatics using tools developed by the BBCF (bbcf.epfl.ch). R.F. and D.N. acknowledge collaborative funding from the Agence Nationale de la Recherche (project “IMP-REGULOME”, ANR-18-CE12-0022-02). R.F. acknowledges funding from the Fondation pour la Recherche Médicale (FRM, DEQ20150331703), the ANR (ANR-13-BSV2-0014) and Labex ‘Epigenmed’, an ANR ‘Investissement d’avenir’ programme (ANR-10-LABX-12-01). D.N. acknowledges funding from the Fondation ARC (PJA20141201727), the FRM (AJE20140630069), the ANR (ANR-14-ACHN-0009-01) and the Fondation Bettencourt-Schueller.

## AUTHOR CONTRIBUTIONS

Conceptualization: D.L., R.F., D.N.; Methodology: D.L., V.P., V.T., R.F., D.N.; Investigation: D.L., B.M., R.P., M.M., B.P., A. M., M.P., A.P., D.N.; Analysis: D.L., B.M., R.P., V.P., B.P., A.M., A.P., R.F., D.N.; Writing: D. L., B.M., R.F., D.N.; Funding acquisition: R.F., D.N.

## Data deposition

Unprocessed ChIP-seq, RNA-seq and 4C-seq sequences are available from the European Nucleotide Archive (EBI-ENA) repository under accession number PRJEB28762. Processed 4C-seq interaction patterns are available from the Mendeley Data repository (http://dx.doi.org/10.17632/6t33n6nm96.2).

## MATERIAL AND METHODS

### ES cells, cell culture and *in vitro* differentiation

Hybrid ESC lines BJ1 ((C57BL/6J x JF1)F1) and JB1 ((JF1 × C57BL/6J)F1) are both male and were derived previously in serum-free (2i) medium (22). These WT cells and the JB1- and BJ1-derived ESC lines with bi-allelic deletions comprising site 2 (this study), and the mono-parental ESC lines PR8 (39) and AK2 (40) were maintained without feeders on gelatin-coated dishes in serum-free ESGRO Complete PLUS medium (Millipore, with LIF and Gsk3 inhibitor). Differentiation of ESCs into cortical neural progenitor cells (NPCs) was performed as described in detail before (22, 41). Briefly, ES cells were plated on matrigel-coated dishes at a density of 3×10^5^ cells per 10-cm dish in serum-free ESGRO Complete PLUS medium and after 24h, the medium was changed to DDM (DMEM/F12 + GlutaMAX (ThermoFisher), supplemented with 1x N2 (ThermoFisher), B27 (without vitamin A, ThermoFisher), 1 mM of sodium pyruvate, 500 ug/ml BSA, 0.1mM of 2-mercapto-ethanol for a total of 12 days. Cyclopamine (1 μM, Merck) was added from day 2 to day 10 of differentiation. Media was changed every two days. After 12 days of differentiation, NPCs were dissociated using StemPro Accutase, and were used for high throughput 4C studies. Part of the cells were replated on poly-lysine (Sigma)/laminin (Sigma) for expression studies, and cultured in 1:1 mixture of DDM and Neurobasal/N27 media (ThermoFisher, supplemented with 1x B27) and 2mM GlutaMax) for further differentiation, till D21, or were re-plated onto coated coverslips to perform immunostainings and DNA-FISH studies two days later (D12+2).

### CRISPR-Cas9 mediated deletion of CTCF binding site 2 at *Meg3* intron1

The guide RNA (sgRNA) was designed using CRISPR Design tool (http://crispr.mit.edu/) and synthesised with *BbsI* sticky ends: *Meg3* DMR CTCF site 2: GTTGCACATAGAGACCGCTAG. It was cloned into the pUC57-sgRNA expression vector (42) (a gift from Xingxu Huang; #51132, Addgene). The Cas9-VP12 vector (43) (a gift from Keith Joung; #72247, Addgene) was modified by adding T2A-GFP at the C-terminal end and electroporated with the sgRNA vector into JB1 and BJ1 hybrid ES cells using the Amaxa nucleofector procedure (Lonza). 24 h post-electroporation, GFP-positive cells were sorted by flow cytometry (FACS Aria, Becton Dickinson) and single cells were seeded onto 96-well plates. After 10-12 days of culture, individual colonies were picked and grown in 6-well plates. Genomic DNA was extracted and the region around *Meg3* DMR CTCF site 2 was amplified (primers in table S4), followed by confirmation of the deletion by DNA sequencing (fig. S6A).

### ChIP-qPCR, ChIP-seq and data analysis

ChIP experiments were performed as previously described (44) with minor modifications. ESCs were fixed for 5 min in a 2% formaldehyde solution at room temperature. ChIP-seq samples were fragmented using a water bath sonicator (BioRuptor Plus, Diagenode). 10 μg of chromatin was immuno-precipitated with either 5 μg of CTCF antibody (07-729, Merck Millipore) or 4 μg of H3K27me3 antibody (17-622, Merck Millipore).

PR8, AK2, BJ1 and JB1 ChIP-qPCR samples were analysed on a LightCycler 480 instrument (Roche Molecular Diagnostics) using the SsoAdvanced Universal SYBR Green Supermix (BioRad). Duplicate qPCR experiments were performed on technical replicates with recovery in each cell type expressed versus a corresponding input sample (primers in table S4).

Indexed ChIP-seq libraries were constructed using the Next Ultra Library Prep Kit for Illumina (New England Biolabs) using the application note ‘Low input ChIP-seq’. Multiplexed sequencing was done using 86-bp single-end reads on the Next-Seq 500 system (Illumina) at the I2BC Next Generation Sequencing Core Facility. Data were mapped to ENSEMBL Mouse assembly GRCm38 (mm10) using BWA with default parameters. Reads for mono-parental PR8 and AK2 samples were further extended to 200 bp. After removal of duplicate, multiple aligning and low-quality reads, densities in windows of 50 bp were calculated for combined technical replicates. Samples were normalised using quantile normalisation after removal of regions with abnormal alignment in the input samples (either ≥3 IQR over median input signal or regions with no input signal at all). CTCF peaks in PR8 and AK2 ESCs were called if four consecutive 50 bp bins had a minimum value of 20 in at least one cell type, followed by extension of one bin left and right (table S1). Differential CTCF peaks were called if the difference between peak values was ≥ 3 fold. We validated differential CTCF binding using ChIP-seq data from JB1 cells. After mapping to ENSEMBL Mouse assembly GRCm38 (mm10) using BWA we identified known JF1 polymorphisms in the reads covering our identified CTCF peaks (ftp://molossinus.lab.nig.ac.jp/pub/msmdb/For_Seq_Analysis/list_of_variations/).

### RNA-seq and data analysis

Total RNA was extracted from AK2 and PR8 ESCs by lysing the cells on the culture dish with the addition of TRI reagent (Sigma-Aldrich). RNA samples were sequenced on the Illumina HiSeq-2000 system (Illumina) using the TruSeqTM SBS stranded mRNA sample kit (version 3). For both ES lines, RNA-sequencing (2× 100bp) was performed in triplicate.

Paired-end Fastq files were mapped to ENSEMBL Mouse assembly GRCm38 (mm10) using STAR (45). Transcript abundance was quantified using RSEM in TPM (Transcript per million) for the 51789 transcripts in EMSEMBL database version 89. Replicates showed high correlation (R ≥ 0.99) indicating good reproducibility and reliability. Samples were normalised against each other using quantile normalisation, followed by averaging of triplicate samples (table S2). Genes were considered significantly detected if the TPM value in either of the combined mono-parental PR8 or AK2 data sets was ≥ 5. Differential expression was called if the highest TPM value was ≥ 5, the fold difference between PR8 and AK2 TPM was ≥ 1.5 and the highest TPM value × the fold TPM difference was ≥ 50.

### RT-qPCR and allele-specific quantitation

Total RNA was extracted from hybrid ESCs and NPCs using the miRNEasy Kit (Qiagen) and DNaseI treatment (Qiagen). cDNA was synthesized from 5μg of RNA using random hexamers and SuperScript III (ThermoFisher) reverse transcriptase. *Meg3* and *Dlk1* expression were quantified by RT-qPCR using SYBR Green I master mix (Roche) on a Lightcycler 480 instrument. Mean C_T_ values were normalized with the mean of two housekeeping genes (*Actb, Gapdh*) and the ΔΔCt method (46). Primer sequences in table S4.

The Taqman mutation detection assay was used for allele-specific quantitation (ThermoFisher). Expression was quantified using Taqman Genotyping Master Mix (ThermoFisher) and a Lightcycler 480 instrument. The levels were normalized to two housekeeping genes (*Actb, Gapdh*) amplified with SYBR Green Master Mix (Roche), as reported before (22).

### Reanalysis of Hi-C data

Raw data for ESCs and NPCs were obtained from GEO dataset GSE96107 (10). HiC-Pro v2.9.0, using Bowtie2 v2.3.0, was used to map the raw data to mouse reference genome mm10 and to process the aligned reads, with default settings to remove duplicates, assign reads to DpnII restriction fragments and filter for valid interactions (47, 48). Binned interaction matrices were generated at 10-kb resolution from the valid interactions and were normalised using the Iterative Correction and Eigenvector decomposition method (ICE) implemented in HiC-Pro. TAD borders were called using TADtool (20), with window size 500 kb and insulation index cut-off value 21.75, resulting in a high degree of genome-wide overlap with TAD borders as reported by Bonev *et al*.

### 4C-seq and data analysis

Chromatin fixation, cell lysis and 4C library preparation were done as previously described (49) using 15 million cells per experiment, *DpnII* (New England Biolabs) as the primary restriction enzyme and *NlaIII* (New England Biolabs) as the secondary restriction enzyme. For 4C-seq library preparation, 800 ng of 4C library was amplified using 16 individual PCR reactions with inverse primers including the Solexa or TruSeq adapter sequences (primers in table S4). Illumina sequencing was done on samples containing PCR amplified material of up to ten viewpoints using 100 bp or 86 bp single end reads on the Illumina Hi-Seq 2500 or Next-Seq 500 systems at the iGE3 Genomics Platform of the University of Geneva (Switzerland) or the I2BC Next Generation Sequencing Core Facility. 4C-seq data sets were mapped and translated into restriction fragments using the 4C-seq pipeline of the BBCF HTSstation (50), according to ENSEMBL Mouse assembly GRCm38 (mm10). For visualisation of 4C-seq patterns, smoothed 4C-seq data (11 fragments) were normalised to the signal within the region covering the 5 TADs surrounding the viewpoint, as described previously (51) (see also fig. S2, C and D). Regions for normalisation: *Igf2-H19* locus – chr7:141,530,000-143,520,000; *Dlk1-Dio3* locus – chr12:105,970,000-110,860,000. Ratios between smoothed 4C-seq patterns were calculated using the BioScript library of the BBCF HTS station (50). Distributions of 4C-seq signal were calculated using a previously described approach (51), with unprocessed 4C-seq data normalised within the previously mentioned 5 TADs and signal within each sub-domain expressed per Mb. Significance of differences in 4C-seq signal on the maternal and paternal chromosomes for individual sub-TADs or between sub-TADs was calculated by determining the fraction of fragments with increased maternal versus paternal signal in sub-domains versus de remainder of the TAD or between sub-domains, followed by a G-test of independence.

### DNA methylation analysis

DNA methylation was analysed by digestion of genomic DNA samples with a methylation sensitive restriction endonuclease, followed by qPCR amplification and Sanger sequencing of the PCR products (22). Briefly, 1 ug of genomic DNA was fragmented with *EcoRI*, after which half of the reaction was further digested with the methylation-sensitive enzyme *Aci*I. 1 ng of both the *Aci*I-digested DNA and of the non-digested (EcoRI only) DNA was used for qPCR. Quantitative values were obtained using the standard curve method (52) with normalisation against two regions without *Aci*I sites (*Col1a2* and *Col9a2*). The *ActB* gene (unmethylated *Aci*I site) and IAP retroposons (methylated *Aci*I sites) were included as controls. Primer sequences in Table S4.

### Probes for 3D DNA-FISH and RNA-FISH

Fosmid and BAC probes were directly labelled by nick translation (Abbott molecular, ref 07J00-001) with Cy3- or Cy5-dUTP (GE Healthcare). Details on fosmids and BACs are provided in table S4. Per coverslip, 0.1 μg of nick-translation product was precipitated in the presence of 10 μg of salmon sperm and 5μg of Cot-1 DNA and resuspended in 10 μl of hybridisation buffer (50% formamide, 2X SSC, 10% dextran sulphate, 1mg/ml BSA, 20mM VRC; pH 7.0) and denatured for 7 min at 75°C. Competition was done for 30 min at 37°C (this step was not applied for RNA-FISH) before overnight hybridisation of the cells.

### 3D DNA-FISH

3D DNA-FISH was carried out on ESCs adhered to 0.1% gelatin-coated coverslips as previously described (22). Briefly, cells were fixed in (3% paraformaldehyde, 1xPBS; pH 7.4) for 10 min at RT and permeabilised for 7 min with (0.5% Triton, 1xPBS; pH 7.4) on ice, and denatured at 80°C in (50% formamide, 2xSSC; pH 7.0) for 30 min. Cells were rinsed in ice-cold 2xSSC; pH 7.0, and hybridised with probes overnight at 42°C (coverslips were sealed onto slides with rubber cement). The next day, cells were washed 3 times in (50% formamide, 2xSSC; pH 7.2) at 42°C, and 3 times in 2xSSC; pH 7.0 at 42°C for 5 min each. Finally, coverslips were stained with DAPI and mounted using Vectashield antifade mounting medium (VectorLabs, H-1000).

### DNA-RNA FISH with MS2 oligo probes

Fixation of hybrid BJ1 ESCs containing 64 copies of MS2 repeats into exon-10 of the Meg3 lncRNA was done as previously described (25). Following fixation, cells were incubated twice for 5 min with DEPC-treated 1xPBS; pH 7.4 at RT and dehydrated in 80%, 95%, 100% ethanol, for 3 minutes each respectively, and were air dried. Then, cells were rehydrated in (20% formamide, 2xSSC, 0.01% Tween 20; pH 7.0) for 5 min at 37°C. An MS2-multi-oligonucleotide probe was mixed in 10μl of hybridisation buffer (20% formamide, 2xSSC, 10% Dextran sulfate, 50mM Sodium phosphate, 2 mM VRC; pH 7.0) that was pre-warmed at 37°C. Probes were denaturated 1 min at 80°C and cells were hybridised with probes for 2h at 37°C in a dark and humid chamber. Coverslips were washed 3 times with (20% formamide, 2xSSC, 0.01% Tween 20; pH 7.0) for 5 min at 37°C, and once briefly in DEPC-treated 1xPBS; pH 7.4. Cells were post-fixed in 4% paraformaldehyde, 1xPBS; pH 7.4 for 10 min at RT and rinsed three times in DEPC-treated 1xPBS; pH 7.4 for 5 min. Cells were incubated in 2xSSC; pH 7.0 for 5 min at 40°C, following denaturation in (70% Formamide, 2xSSC, 50mM Sodium Phosphate buffer; pH 7.0) for 3 min at 73°C, and then in (50% Formamide, 2xSSC, 50mM Sodium Phosphate buffer; pH 7.0) for 1 min at 73°C. Cells were hybridised with prepared fosmid probes overnight at 37°C (coverslips were sealed onto slides with rubber cement). The next day, cells were washed 3 times in (50% formamide, 2xSSC, 0.01% Tween 20; pH 7.2) at 42°C for 5 min each, and 3 times in (2xSSC; pH 7.0) at 42°C for 5 min each. Finally, coverslips were stained with DAPI and mounted using Vectashield antifade mounting medium.

### Immunofluorescence

Immunofluorescence staining was performed as described (41). Primary antibodies: anti-Nestin (839801, Biolegend; 1:1000 dilution), anti-Tubb3 (801201, Biolegend; 1:1000 dilution). Secondary antibodies: goat anti-mouse Alexa Fluor 488 (A-11011, Life Technologies; 1:1000 dilution), goat antirabbit Alexa Fluor 594 (A-11012, Life Technologies; 1:1000 dilution).

### Confocal Microscopy and data analysis

Three-dimensional images were acquired with a Zeiss LSM780 laser scanning confocal microscope (Zeiss), using a 63xNA 1.4 Plan-Apochromat oil immersion objective. Optical sections separated by 0.4 μm steps were collected in the Z direction. Stacks were analysed using Imaris software (Bitplane, Switzerland). FISH signals were segmented in 3D and their centres of mass were defined. For double FISH experiments, the distances between closest neighbour’s centre of mass were calculated. Only FISH fluorescence signals within DAPI 3D-segmented object were considered for the analysis.

Measured distances between BAC probe FISH signals were normalised to the genomic distances and not to the radius of individual nuclei, because no differences in cell radius were observed between the three cells lines (PR8, AK2 and BJ1) (not shown). Significance of differences between distances from combined repeated experiments of DNA probes were calculated using the two-tailed unpaired Mann-Whitney t-test.

---

**Figure S1.**
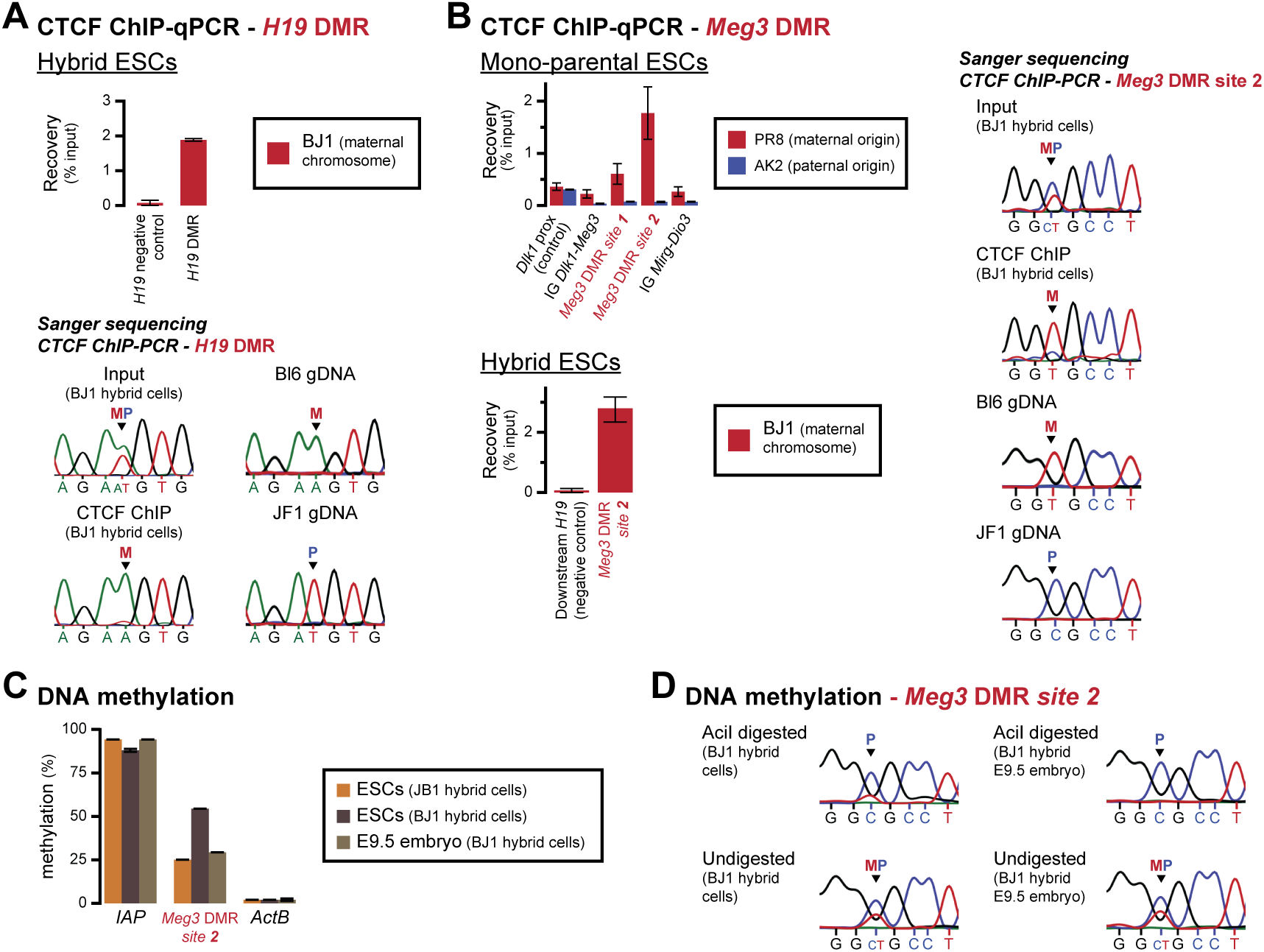
Multiple instances of bi-allelic CTCF binding accompany maternal allele-specific CTCF binding at the *Meg3* and *H19* DMRs. **A.** ChIP-qPCR (top) and ChIP-PCR followed by Sanger sequencing (bottom) confirms maternal allele-specific CTCF binding at the *H19* DMR in hybrid ESCs. Error bars indicate SE from 2 replicates. **B.** Top left: ChIP-qPCR validation of maternal allele-specific CTCF binding at the *Dlk1-Dio3* locus in mono-parental PR8 and AK2 ESCs. Error bars indicate SE from 2 replicates. Bottom left: ChIP-qPCR validation of maternal allele-specific CTCF binding at site 2 in the *Meg3* DMR in hybrid ESCs. Error bars indicate SE from 2 replicates. Right: ChIP-PCR followed by Sanger sequencing (bottom) confirms maternal allele-specific CTCF binding at site 2 in the *Meg3* DMR in hybrid ESCs. **C.** DNA methylation levels as determined by digestion of genomic DNA from hybrid ESCs and E9.5 hybrid embryos with *Aci*I, an endonuclease that cuts non-methylated DNA only. Values are expressed as percentage of non-digested DNA. Positive control: IAP transposable elements (high levels of methylation) and negative control: *ActB* promoter (low levels of methylation). Methylation of CTCF site 2 in the *Meg3* DMR in ESCs is in a similar range as in E9.5 embryo. **D.** Confirmation of maternal-specific DNA methylation in hybrid ESCs and E9.5 embryos by Sanger sequencing of genomic DNA with (top) and without (bottom) methylation-sensitive *Aci*I digestion. Parental origin of the SNP that distinguishes the maternal and paternal alleles is indicated. See also Fig. 1D.

**Figure S2.**
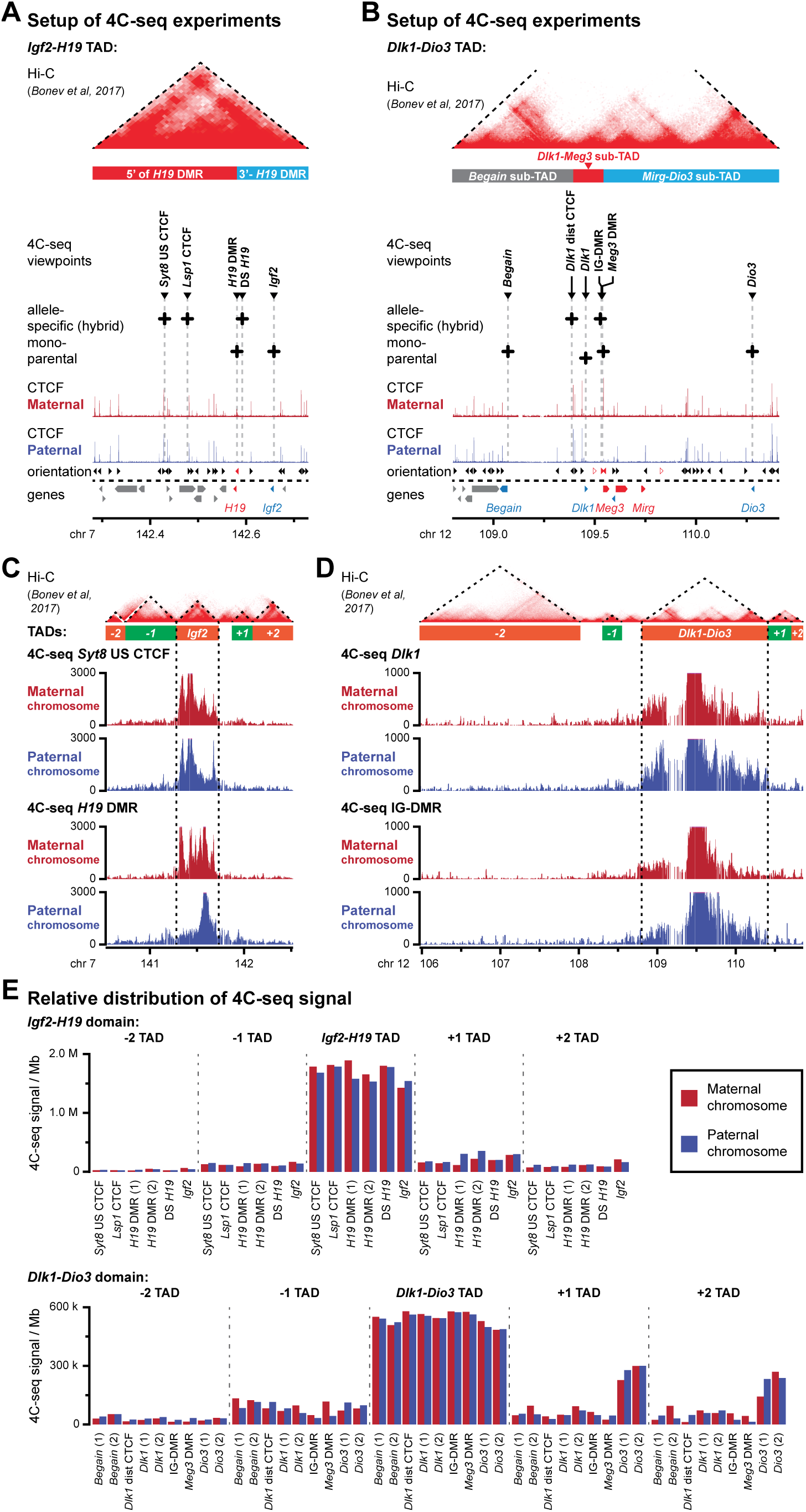
DNA interactions at paternally imprinted gene domains are restricted within the same TADs on both the parental chromosomes. **A.** Set-up of 4C-seq experiments at the paternally imprinted *Igf2-H19* domain and its surrounding TAD. The positions of viewpoints in hybrid cells (allele-specific) or mono-parental cells are indicated within the overarching TAD. Allele-specific CTCF signal is indicated below. Non-allelic Hi-C signal and the position of the maternal sub-TADs are indicated above. The orientation of CTCF sites and genes are indicated below with colours indicating allele-specificity. **B.** Set-up of 4C-seq experiments at the paternally imprinted *Dlk1-Dio3* domain and its surrounding TAD. **C.** 4C-seq signal for two viewpoints in the *Igf2-H19* domain on the maternal (red) and paternal (blue) chromosome in mono-parental and hybrid ESCs. Signal is indicated in the 5 TADs surrounding the viewpoint (orange-green blocks), with reanalysed Hi-C signal visualized above. Allele-specific CTCF signal (ChIP-seq on mono-parental ESCs) is provided below. Hi-C data are from reference (10). **D.** 4C-seq signal for two viewpoints in the *Dlk1-Dio3* domain on the maternal (red) and paternal (blue) chromosome in mono-parental and hybrid ESCs. **E.** Relative distribution of 4C-seq signal for indicated viewpoints in the 5 TADs surrounding the two imprinted domains.

**Figure S3.**
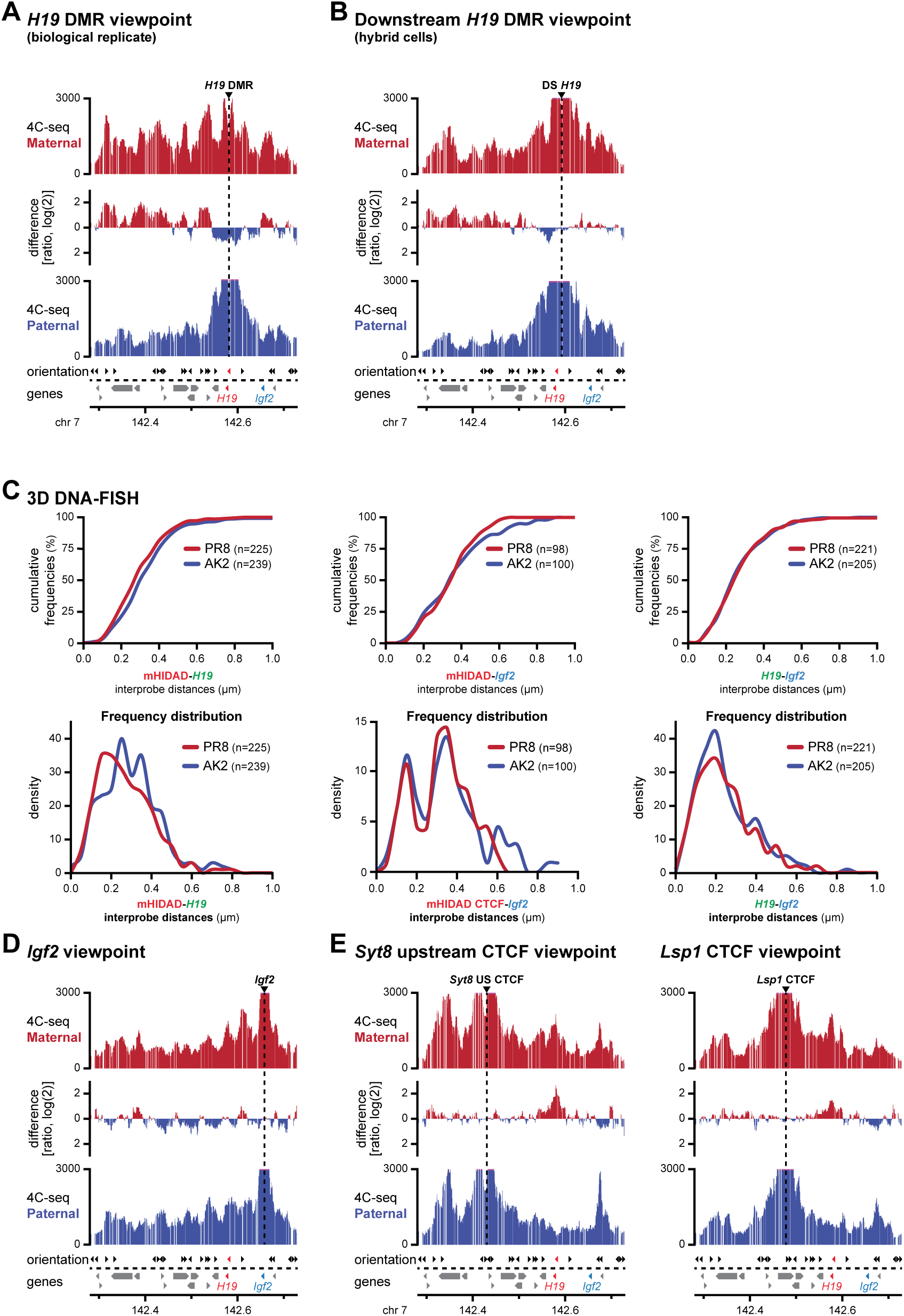
The *Igf2-H19* domain adopts an allele-specific sub-TAD organisation that is anchored by bi-allelic and allele-specific CTCF sites. **A.** 4C-seq signal from a biological replicate for the *H19* DMR viewpoint on the maternal (red) and paternal (blue) chromosomes in mono-parental ESCs. The ratio of interactions is provided between the patterns. **B.** 4C-seq signal for a viewpoint 10 kb downstream of the *H19* DMR in hybrid ESCs. **C.** Distance distribution and cumulative distance frequencies between indicated fosmid probes in mono-parental ESCs. **D.** 4C-seq signal for the *Igf2* viewpoint in mono-parental ESCs. **E.** 4C-seq signal for two bi-allelic CTCF peaks (left: CTCF peak upstream of the *Syt8* gene and right: CTCF peak in the *Lsp1* gene) in hybrid ESCs.

**Figure S4.**
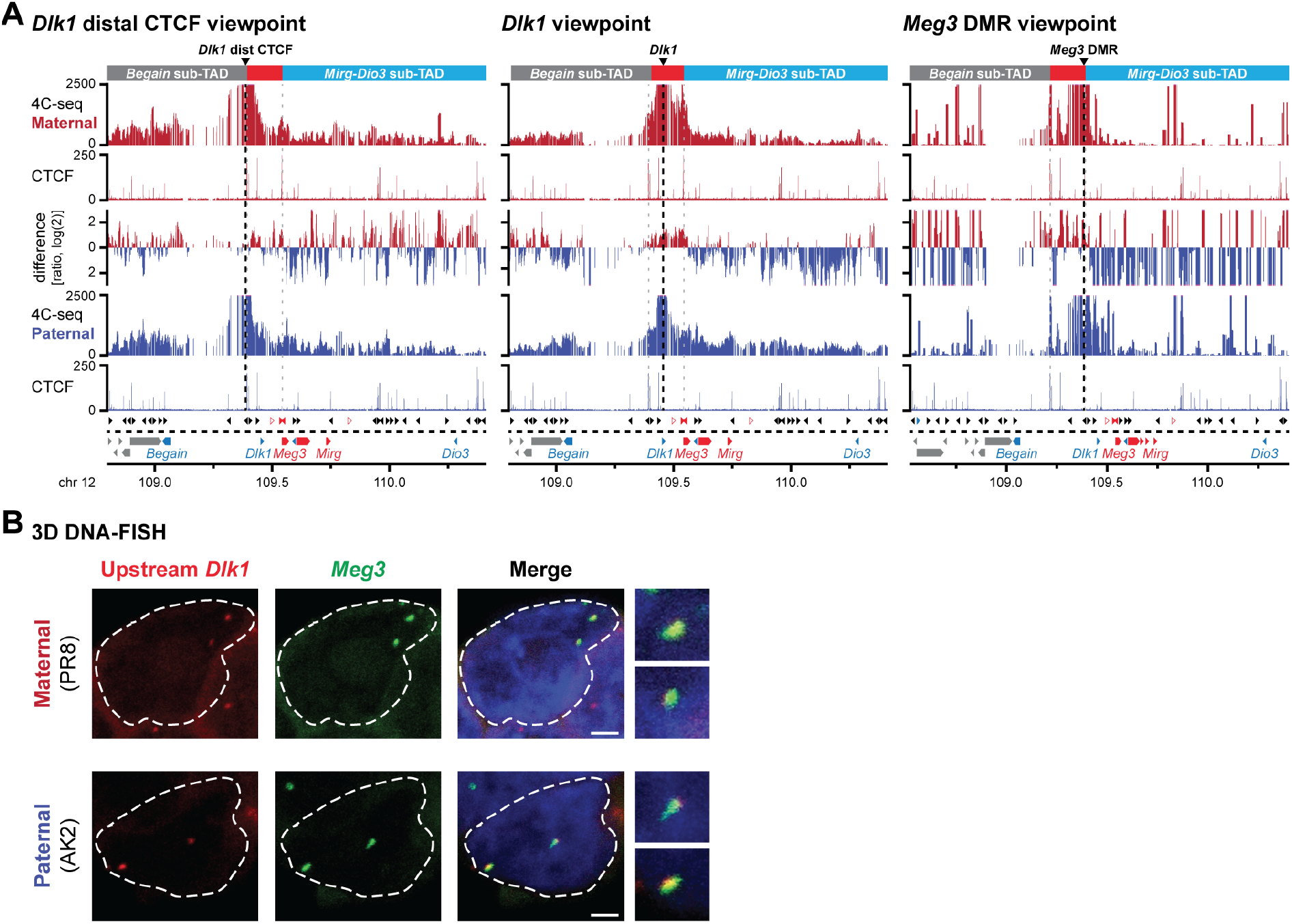
A maternal *Dlk1-Meg3* sub-TAD is structured by bi-allelic and allele-specific CTCF sites. **A.** 4C-seq signal for indicated viewpoints in the *Dlk1-Meg3* sub-TAD on the maternal (red) and paternal (blue) alleles in the 1.6-Mb *Dlk1-Dio3* TAD. The ratio of interactions is provided in-between the patterns. Allele-specific CTCF signal is indicated below each 4C pattern. 4C-seq viewpoints and sub-TADs are indicated above. The orientation of CTCF sites and genes are indicated below with colours indicating allele-specificity. **B.** Representative examples of DNA-FISH using fosmid probes in mono-parental ESCs. Scale bar, 2 μm. See also Fig. 3C.

**Figure S5.**
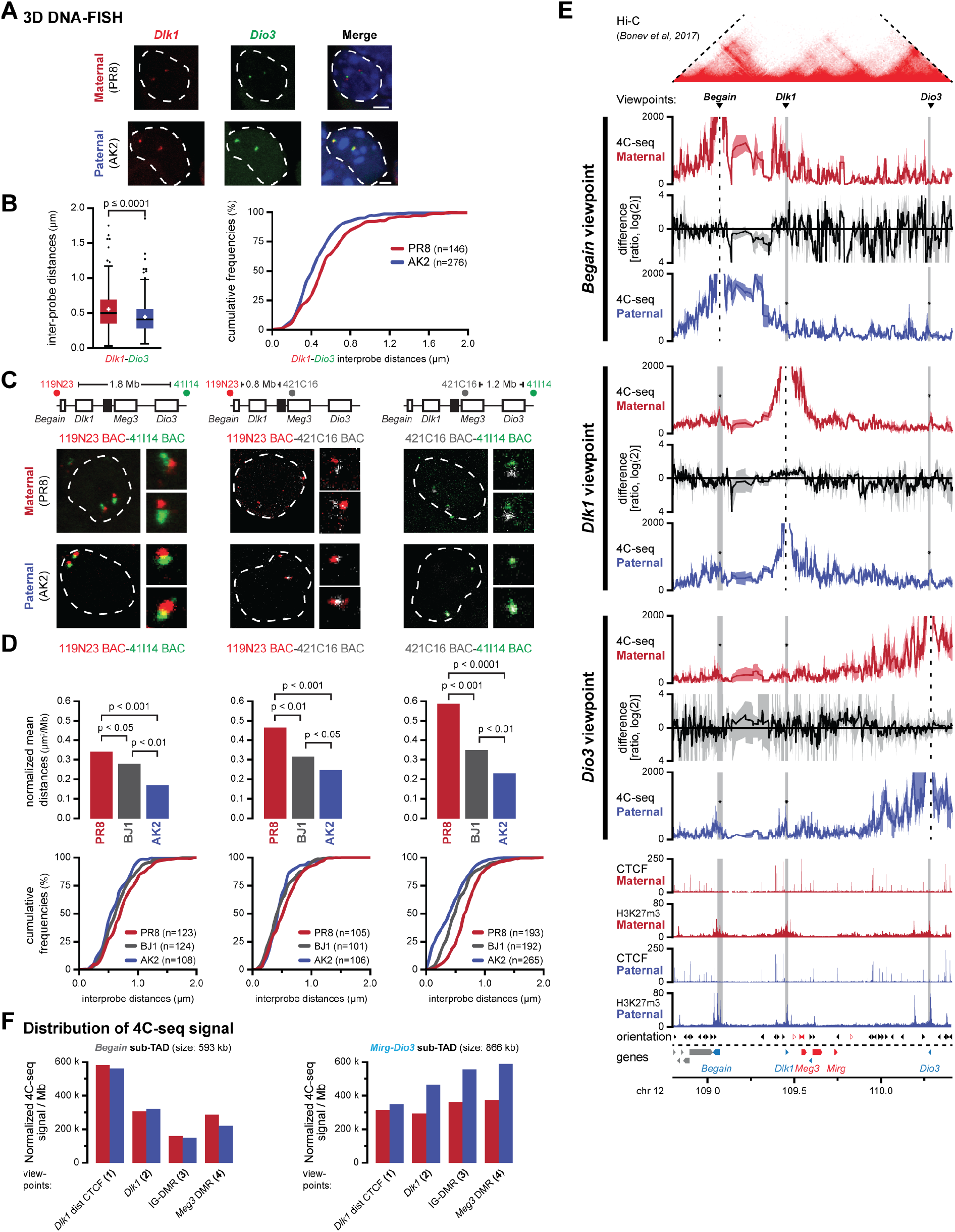
The *Dlk1-Dio3* domain is organised into allele-specific sub-TADs that coincide with different allelic intra-TAD distances. **A.** Examples of 3D DNA-FISH with fosmid probes in the *Dlk1-Dio3* TAD. Images show representative cells in mono-parental ESCs. Scale bars, 2 μm. **B.** Distance measurements in mono-parental cells confirm the increased separation between *Dlk1* and *Dio3* on the maternal chromosome. **C.** Representative examples of DNA-FISH using three BAC probes in mono-parental and hybrid ESCs. Scale bars, 1 μm. The schematic location of the BAC probes is indicated above. **D.** Distance measurements in mono-parental and hybrid cells confirm the increased separation between all combinations of BAC probes on the maternal chromosome. **E.** 4C-seq line-graphs for the inactive, H3K27me3-marked protein-coding genes at the *Dlk1-Dio3* locus. Lines indicate the average 4C-seq signal from 2-3 replicates/viewpoint, with surface showing maximum and minimum values. The ratio of interactions between replicate experiments is provided between the patterns, with the surface showing maximum difference between samples. The *Dlk1-Dio3* TAD (Hi-C signal) and the 4C-seq viewpoints are indicated above. Allele-specific CTCF and H3K27me3 signal (ChIP-seq), the orientation of CTCF sites and genes are indicated below. A moderate enrichment of interactions at or near the H3K27me3-marked promoters of the imprinted protein-coding genes within the locus may be observed (asterisks), yet with little difference between the maternal and paternal alleles. **F.** Distribution of 4C-seq signal for indicated viewpoints in the *Begain* sub-TAD (left) and the *Mirg-Dio3* sub-TAD (right). Whereas interactions in the *Begain* sub-TAD are largely invariant between the parental chromosomes, in the *Mirg-Dio3* sub-TAD they are consistently enriched on the paternal chromosome.

**Figure S6.**
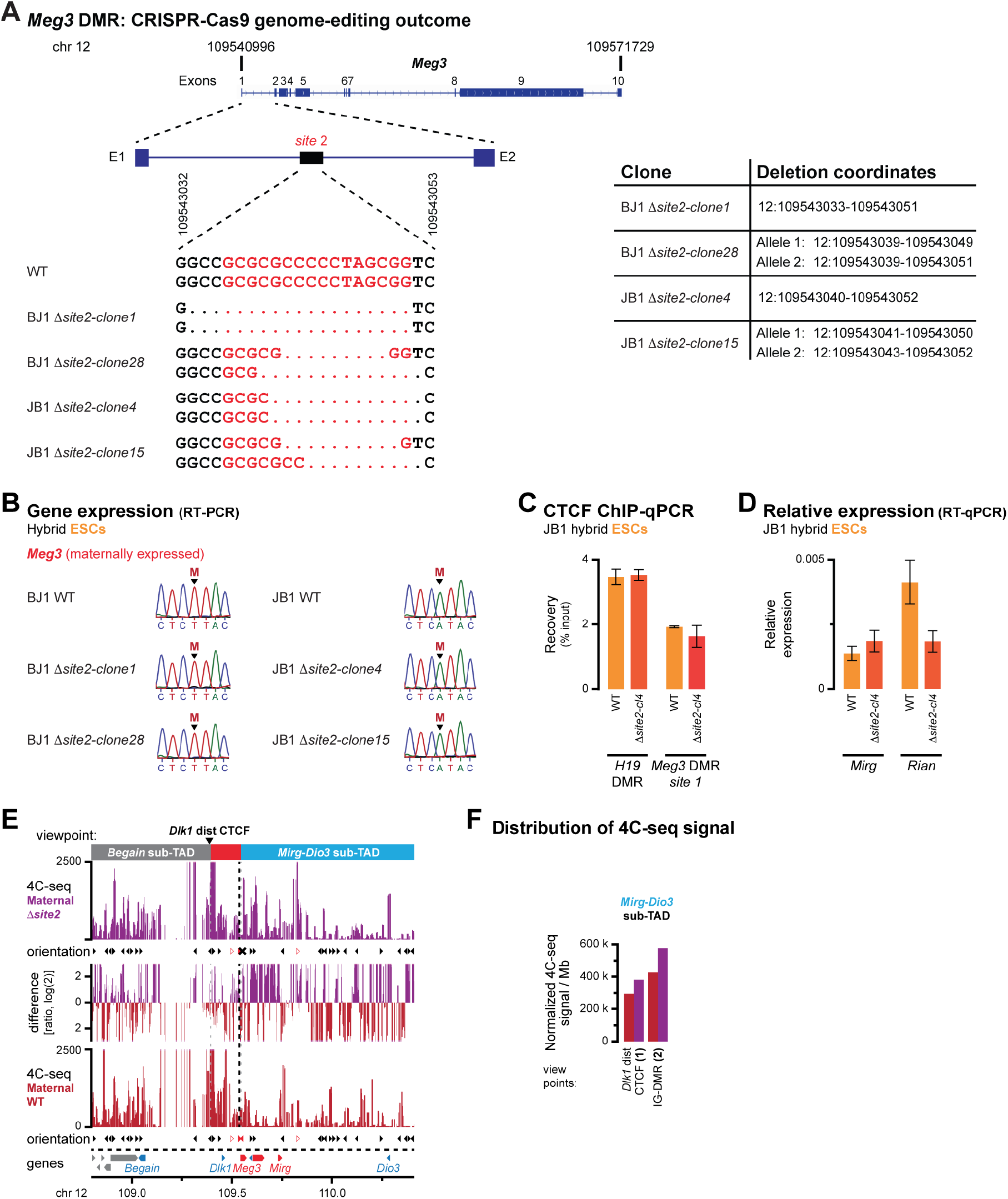
CTCF binding at site 2 in the *Meg3* DMR is required for the structure of the *Dlk1-Meg3* sub-TAD. **A.** Location of CTCF binding site 2 in the *Meg3* gene (top) and genotyping of the CTCF binding site deletions in the hybrid BJ1 and JB1 clones generated in this study (bottom). Coordinates of deletions (mm10) are provided in the panel on the right. **B.** Sanger sequencing of RT-PCR products confirms continued maternal allele-specific *Meg3* expression in JB1- and BJ1-derived hybrid ESC lines carrying a deletion in CTCF binding site 2 in the *Meg3* DMR. The parental origin of the SNP that distinguishes the maternal and paternal alleles is indicated. **C.** ChIP-qPCR validation of CTCF binding at site 1 in the *Meg3* DMR in hybrid ESCs. Error bars indicate SE from 2 replicates. **D.** Expression levels of *Mirg* and *Rian* non-coding RNAs in ESCs with a deleted CTCF site 2 in the *Meg3* DMR or their WT counterparts. **E.** 4C-seq signal for the distal *Dlk1* CTCF peak on the maternal alleles from ESCs with a deleted CTCF site 2 in the *Meg3* DMR (purple) or their WT counterparts (red) in the entire *Dlk1-Dio3* TAD. The ratio of interactions is provided in-between. The orientation of CTCF sites is indicated below each panel, with an X indicating the deleted CTCF site. The position of the viewpoint and the sub-TADs are indicated above (red box: *Dlk1-Meg3* sub-TAD). **F.** Distribution of 4C-seq signal for indicated viewpoints in the *Mirg-Dio3* sub-TAD. In the deletion cells, 3D interactions are reorganized similar to the paternal allele, with increased 4C-seq signal in the *Mirg-Dio3* sub-TAD.

**Figure S7.**
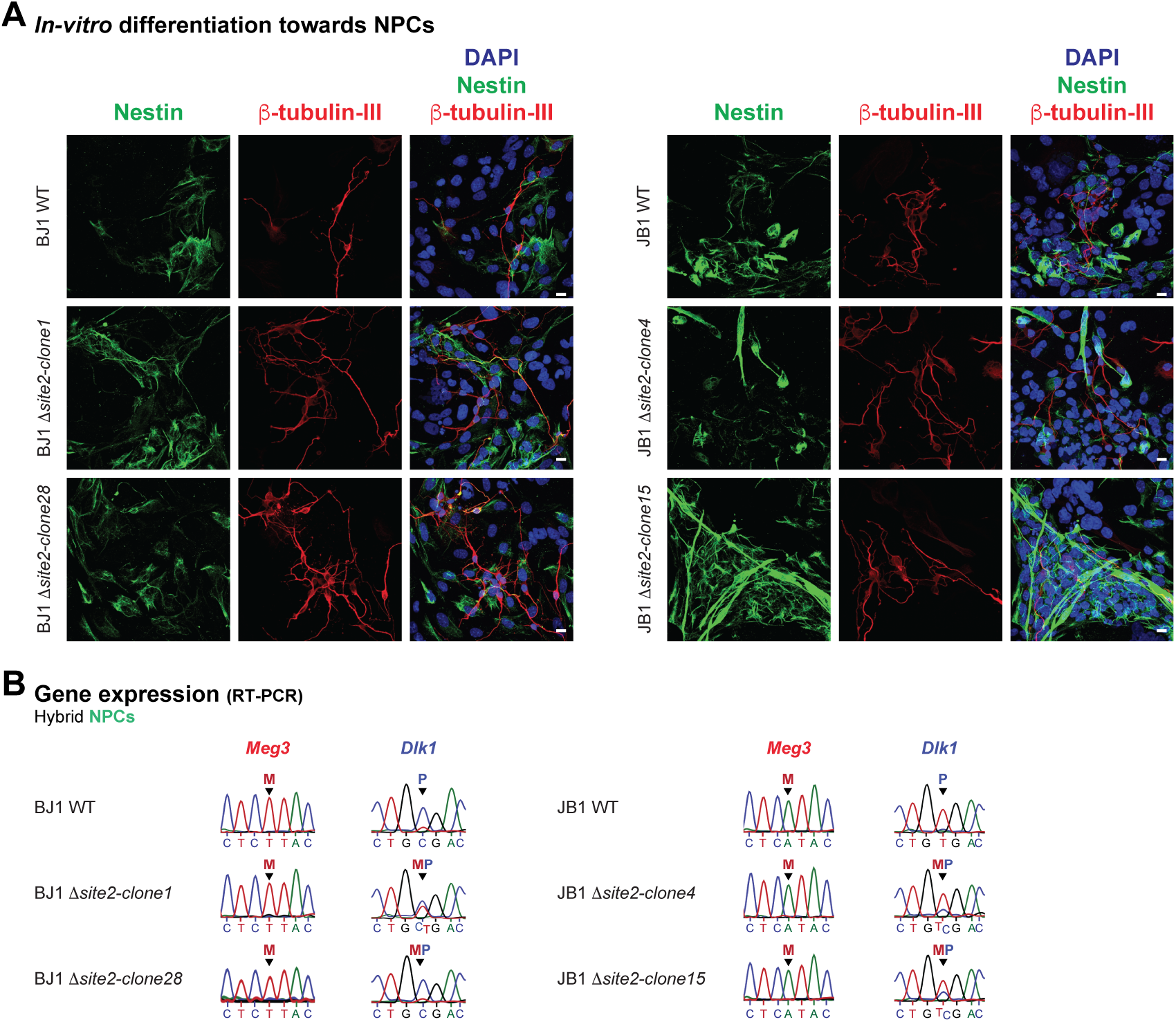
CTCF binding at site 2 in the *Meg3* DMR is required for correct imprinted activation of *Dlk1*. **A.** Immunofluorescence staining of Nestin (neuronal progenitors) and Tubulin-β3 (neurons) in NPCs with a deleted CTCF site 2 in the *Meg3* DMR or their WT counterparts. Cells were counterstained with DAPI. Scale bars, 10 μm. **B.** Maternal allele-specific *Meg3* expression and bi-allelic *Dlk1* activation in *in vitro* differentiated NPCs with a deleted CTCF site 2 in the *Meg3* DMR or their WT counterparts. Determination of allele specificity by Sanger sequencing of RT-PCR products. Parental origin of the SNP that distinguishes the maternal and paternal alleles is indicated.

**Figure S8.**
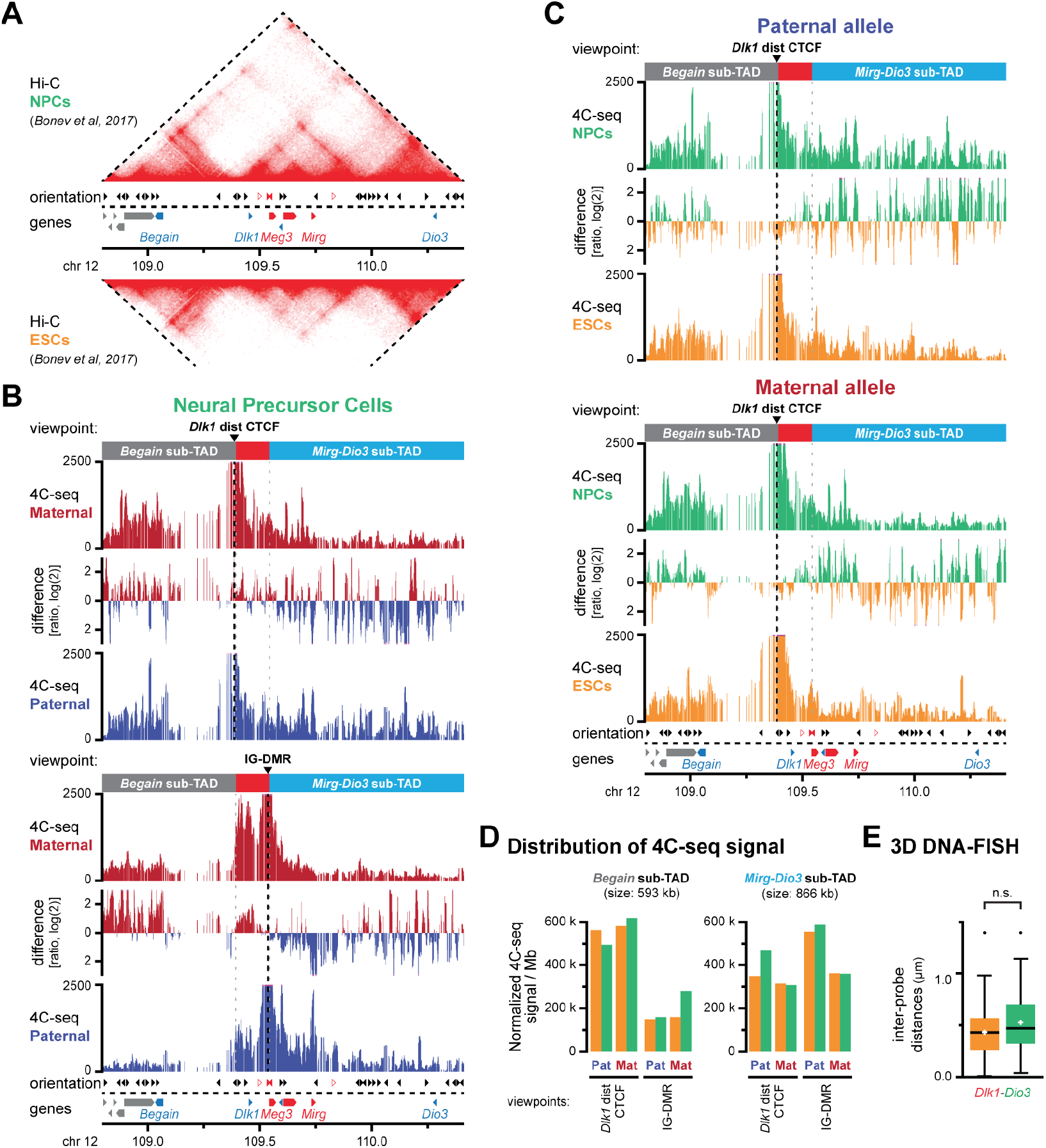
Developmental *Dlk1* activation on the maternal chromosome coincides with a mostly stable intra-TAD organisation. **A.** Reanalysed non-allelic Hi-C signal from NPCs (top) and ESCs (bottom) in the *Dlk1-Dio3* locus. The orientation of CTCF sites identified in ESCs and genes are indicated below with colours indicating allele-specificity. Hi-C data are from reference (10). **B.** 4C-seq signal for the distal *Dlk1* CTCF peak (top) and the IG-DMR (bottom) in NPCs cells on the maternal (red) and paternal (blue) allele in the *Dlk1-Dio3* TAD. The ratio of interactions is provided between the patterns. The position of the 4C-seq viewpoints and the sub-TADs are indicated above (red box: *Dlk1-Meg3* sub-TAD). **C.** 4C-seq signal for the distal *Dlk1* CTCF peak in *in vitro* differentiated hybrid NPCs (green) and ESCs (orange) on the paternal (top) and maternal (bottom) allele in the *Dlk1-Dio3* TAD. The ratio of interactions is provided between the patterns. **D.** Distribution of 4C-seq signal in NPCs and ESCs for indicated viewpoints in the *Begain* and the *Mirg-Dio3* sub-TADs. **E.** 3D DNA-FISH distance measurements with fosmid probes (see Fig. 3F) reveal no significant difference in distances between *Dlk1* and *Dio3* in hybrid NPCs and ESCs.

