## Supplementary material for "CTCF controls imprinted gene activity at the mouse *Dlk1-Dio3* and *Igf2-H19* domains by modulating allele-specific sub-TAD structure": Table S3

### Table S3: Significance of 4C-seq signal in sub-TADs

Significance of 4C-seq signal for individual sub-TADs or differences between sub-TADs was calculated by determining the fraction of fragments with increased maternal versus paternal signal in individual sub-domains versus the remainder of the TAD, or by comparing the score between individual sub-domains. Significance of difference was determined using a G-test of independence.
#: too few instances of signal for reliable calculation of test-statistic.

Color gradient: P < 0.001 | P < 0.01 | P < 0.05

#### *A: Igf2- H19* sub-domains – maternal vs. paternal; ESCs

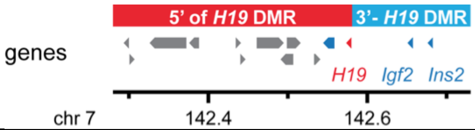


| Viewpoint | P-value<br>5' of H19 DMR *vs.* 3' of H19 DMR |
|---|---|
| *Syt8* upstream CTCF | 0.18 |
| *Lsp1* CTCF | 0.08 |
| *H19* DMR – replicate 1 | $2.90 \times 10^{-7}$ |
| *H19* DMR – replicate 2 | $4.98 \times 10^{-7}$ |
| DS *H19* | 0.18 |
| *Igf2* | 0.23 |

#### *B: Dlk1-Dio3* domain – maternal vs. paternal; ESCs

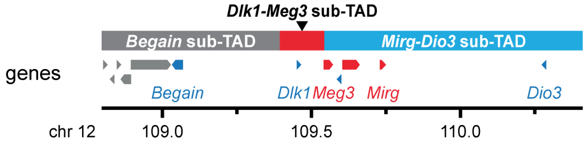


| Viewpoint | P-value | | |
|---|---|---|---|
| | *Begain* sub-TAD *vs.* remainder TAD | *Dlk1-Meg3* sub-TAD *vs.* remainder TAD | *Mirg-Dio3* sub-TAD *vs.* remainder TAD |
| *Begain* – replicate 1 | 0.36 | 0.77 | 0.69 |
| *Begain* – replicate 2 | $2.19 \times 10^{-8}$ | 0.19 | 0.02 |
| *Dlk1* distal CTCF | $1.41 \times 10^{-3}$ | $2.07 \times 10^{-3}$ | 0.08 |
| *Dlk1* – replicate 1 | 0.75 | $2.05 \times 10^{-10}$ | 0.04 |
| *Dlk1* – replicate 2 | 0.52 | $2.72 \times 10^{-8}$ | 0.07 |
| IG-DMR | $6.25 \times 10^{-3}$ | $9.26 \times 10^{-21}$ | $1.11 \times 10^{-4}$ |
| *Meg3* | 0.38 | $3.52 \times 10^{-3}$ | 0.08 |
| Intergenic *Mirg-Dio3* | 0.14 | 0.31 | 0.92 |
| *Dio3* – replicate 1 | 0.07 | # | 0.66 |
| *Dio3* – replicate 2 | 0.11 | 0.13 | 0.43 |
| *Dio3* – replicate 3 | 0.13 | 0.64 | 0.78 |

#### *C: Dlk1-Dio3* sub-domains – maternal vs. paternal; ESCs

| Viewpoint | P-value | |
|---|---|---|
| | *Dlk1-Meg3* sub-TAD *vs.* *Begain* sub-TAD | *Dlk1-Meg3* sub-TAD *vs.* *Mirg-Dio3* sub-TAD |
| *Dlk1* distal CTCF | 0.42 | $4.38 \times 10^{-4}$ |
| *Dlk1* – replicate 1 | $1.06 \times 10^{-6}$ | $1.97 \times 10^{-9}$ |
| *Dlk1* – replicate 2 | $4.35 \times 10^{-5}$ | $5.54 \times 10^{-8}$ |
| IG-DMR | $9.67 \times 10^{-9}$ | $9.27 \times 10^{-21}$ |
| *Meg3* | 0.57 | $5.09 \times 10^{-3}$ |

#### *D: Dlk1-Dio3* domain – maternal vs. paternal; NPCs

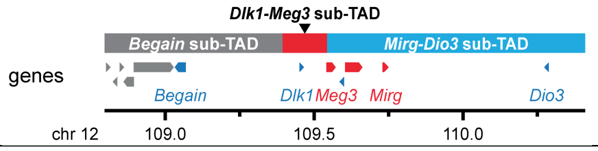


| **Viewpoint** | **P-value** | | |
|---|---|---|---|
| | ***Begain* sub-TAD *vs.* remainder TAD** | ***Dlk1-Meg3* sub-TAD *vs.* remainder TAD** | ***Mirg-Dio3* sub-TAD *vs.* remainder TAD** |
| *Dlk1* distal CTCF | $3.48 \times 10^{-6}$ | $3.14 \times 10^{-5}$ | $5.61 \times 10^{-4}$ |
| IG-DMR | $3.29 \times 10^{-6}$ | $2.84 \times 10^{-24}$ | $6.20 \times 10^{-7}$ |
