## Supplementary material for "CTCF controls imprinted gene activity at the mouse *Dlk1-Dio3* and *Igf2-H19* domains by modulating allele-specific sub-TAD structure": Table S4

### Table S4: Primers and probes

#### *Primer sequences for indicated experiments*

RT-PCR – hybrid cells (*Mus musculus* C57Bl6 genome)

| Gene Name | Sequences (5'-3') | | Source: |
|---|---|---|---|
| *Dlk1* | Fwd: | TTGCTCCTGCTGGCTTTC | Sanli et al., 2018 |
| | Rev: | CCTTGCAGACTCCATTGACA | |
| *Meg3* | Fwd: | CACAGAAGACGAAGAGCTGGA | Sanli et al., 2018 |
| | Rev: | GGTAGAGGTGCACAGCAGGT | |
| *Rian* | Fwd: | CAATGGGTGGATCGTACCTC | Sanli et al., 2018 |
| | Rev: | GTGCTGCCTCAGTCTTTGTG | |
| *Mirg* | Fwd: | TCGGCAGTACATACCAGGTG | Sanli et al., 2018 |
| | Rev: | ACTGATGGCTTCAGGTCAGG | |
| *Gapdh* | Fwd: | CGTCCCGTAGACAAAATGGT | Sanli et al., 2018 |
| | Rev: | TGACTGTGCCGTTGAATTTG | |
| *ActB* | Fwd: | GGCCCAGAGCAAGAGAGGTATCC | Leeb et al., 2010 |
| | Rev: | ACGCACGATTTCCCTCTCAGC | |

ChIP-qPCR – monoparental cells (*Mus musculus* C57Bl6 genome)

| | | |
|---|---|---|
| *Dlk1* prox | Fwd: | CTAGGCGGGGCAGGTGTGCT |
| | Rev: | GAAGGCCCCAGAAGGCTCGC |
| IG *Dlk1-Meg3* | Fwd: | AGCACGCTCGCACTGAACCTG |
| | Rev: | GAAGGGAGGAGCAGAGCCCAGAG |
| *Meg3* DMR (site **1**) | Fwd: | CGCATGATGGCTGCGGCTAGATT |
| | Rev: | AGCCCAGAATGAGGAGGGGGC |
| *Meg3* DMR (site **2**) | Fwd: | CCCCCTCCTACGTCAGTCTAGCTCT |
| | Rev: | AAGACTCCAATAGCCCAACCACCTGAG |
| Intergenic *Mirg-Dio3* | Fwd: | TCCAGCCTGTGCTTGGCTGC |
| | Rev: | GTGTGAGCTGGGGGCGTGTC |

ChIP-qPCR – hybrid cells (*Mus musculus* C57Bl6 genome)

| | | | |
|---|---|---|---|
| *Meg3* DMR (site **1**) | Fwd: | CTTTTGCCGTTTCCTTTGTC | Chatzinikolaou et al., 2017 |
| | Rev: | AACAAAGGCCACCTCCTCTT | |
| *Meg3* DMR (site **2**) | Fwd: | CCCCCTCCTACGTCAGTCTA | |
| | Rev: | AGGTCACAAGTGTTAGCTGTGTG | |
| *H19* DMR | Fwd: | CATGCTTAGTGGGGTCTGCA | Chatzinikolaou et al., 2017 |
| | Rev: | GCCATCAGCGCTATTGTGTG | |
| Downstream *H19* (negative control) | Fwd: | CGCATGGCACCAGAGAAGTA | Chatzinikolaou et al., 2017 |
| | Rev: | TCAGCCCCAAACAGAATCCC | |

4C-seq HiSeq2500 – monoparental cells (*Mus musculus* C57Bl6 genome)

| | | |
|---|---|---|
| *H19* DMR | Fwd: | AATGATACGGCGACCACCGAACACTCTTTCCCTACACGACGCTCTTCCGATCTATTGTTTGCAGCCCTGAGCC |
| | Rev: | CAAGCAGAAGACGGCATACGACTCAGACCCCATAAACCAGTGC |
| *Igf2* | Fwd: | AATGATACGGCGACCACCGAACACTCTTTCCCTACACGACGCTCTTCCGATCTCAGCCTCTGTCCTATGCCCC |
| | Rev: | CAAGCAGAAGACGGCATACGAAACAGCCCCCATACCCCC |
| *Begain* | Fwd: | AATGATACGGCGACCACCGAACACTCTTTCCCTACACGACGCTCTTCCGATCTTCCTGATTACCAAAGACAACAGTCA |
| | Rev: | CAAGCAGAAGACGGCATACGATTAGCACTGGGGAGAGCTGG |
| *Dlk1* | Fwd: | AATGATACGGCGACCACCGAACACTCTTTCCCTACACGACGCTCTTCCGATCTGGCCTTCCTAACCCTCAGCA |
| | Rev: | CAAGCAGAAGACGGCATACGAGCTCTCCTGTCCATACGGGT |
| *Meg3* | Fwd: | AATGATACGGCGACCACCGAACACTCTTTCCCTACACGACGCTCTTCCGATCTAACCTATGCTAATGTTTGGATGGGA |
| | Rev: | CAAGCAGAAGACGGCATACGAGCCAGAAGGACAAACATGTTGC |
| *Dio3* | Fwd: | AATGATACGGCGACCACCGAACACTCTTTCCCTACACGACGCTCTTCCGATCTCCACTGACTGCTTGGCTCTG |
| | Rev: | CAAGCAGAAGACGGCATACGACTACAGCTCCAGCTGCTTGC |

### <u>4C-seq HiSeq2500</u> – hybrid cells (*Mus musculus molossinus* JF1 genome)

| | | |
|---|---|---|
| *Syt8* upstream CTCF | Fwd: | AATGATACGGCGACCACCGAACACTCTTTCCCTACACGACGCTCTTCCGATCTTGGTCACAGCTCTCCAAGTCT |
| | Rev: | CAAGCAGAAGACGGCATACGAGGGGCTAGGGTCTACAGCAA |
| *Lsp1* CTCF | Fwd: | AATGATACGGCGACCACCGAACACTCTTTCCCTACACGACGCTCTTCCGATCTCCCACACCTCATCCAGAGGC |
| | Rev: | CAAGCAGAAGACGGCATACGATGCCTCACCTGAGTGTGCAT |
| DS *H19* | Fwd: | AATGATACGGCGACCACCGAACACTCTTTCCCTACACGACGCTCTTCCGATCTGGGTCCAGAACCCACTTTCTGA |
| | Rev: | CAAGCAGAAGACGGCATACGATTTCCCCAGAGTAGGGGCTG |
| *Dlk1* distal CTCF | Fwd: | AATGATACGGCGACCACCGAACACTCTTTCCCTACACGACGCTCTTCCGATCTACTAACCGGGGTCTCTCACG |
| | Rev: | CAAGCAGAAGACGGCATACGACCTTCAGAACTTTGAGCTAAATAAACCT |
| IG-DMR | Fwd: | AATGATACGGCGACCACCGAACACTCTTTCCCTACACGACGCTCTTCCGATCTTCTTCTATCAGCCCTAAGAATCCTGA |
| | Rev: | CAAGCAGAAGACGGCATACGAATAACCCTGCGGAATGGGTG |

### <u>4C-seq NextSeq500</u> – hybrid cells (*Mus musculus molossinus* JF1 genome)

| | | |
|---|---|---|
| *Dlk1* distal CTCF | Fwd: | AATGATACGGCGACCACCGAGATCTACACTCTTTCCCTACACGACGCTCTTCCGATCTACTAACCGGGGTCTCTCACG |
| | Rev index 1: | CAAGCAGAAGACGGCATACGAGAT**CGTGAT**GTGACTGGAGTTCAGACGTGTGCTCTTCCGATCTCCTTCAGAACTTTGAGCTAAATAAACCT |
| | Rev index 2: | CAAGCAGAAGACGGCATACGAGAT**ACATCG**GTGACTGGAGTTCAGACGTGTGCTCTTCCGATCTCCTTCAGAACTTTGAGCTAAATAAACCT |
| IG-DMR | Fwd: | AATGATACGGCGACCACCGAGATCTACACTCTTTCCCTACACGACGCTCTTCCGATCTTCTTCTATCAGCCCTAAGAATCCTGA |
| | Rev index 1: | CAAGCAGAAGACGGCATACGAGAT**CGTGAT**GTGACTGGAGTTCAGACGTGTGCTCTTCCGATCTATAACCCTGCGGAATGGGTG |
| | Rev index 2: | CAAGCAGAAGACGGCATACGAGAT**ACATCG**GTGACTGGAGTTCAGACGTGTGCTCTTCCGATCTATAACCCTGCGGAATGGGTG |

### <u>Genotyping and ChIP-PCR followed by Sanger sequencing</u>

| | | |
|---|---|---|
| CTCF site in *Meg3* DMR (site **2**) | Fwd: | CCCCCTCCTACGTCAGTCTA |
| | Rev: | AGGTCACAAGTGTTAGCTGTGTG |
| CTCF site in *H19* DMR | Fwd: | GCGCAAATCACCAGACTTTC |
| | Rev: | CAGGTATCTGACTTATAGGGTTCTG |

### <u>DNA methylation studies</u> – hybrid cells

| | | | |
|---|---|---|---|
| *Meg3* DMR (site **2**) | Fwd: | CCCCCTCCTACGTCAGTCTA | |
| | Rev: | AGGTCACAAGTGTTAGCTGTGTG | |
| *ActB* | Fwd: | GGCTTTCCGGCTATTGCTA | |
| | Rev: | CCTCTGGGTGTGGATGTCA | |
| *IAP* | Fwd: | CAAATTAAAGAGCTTGCCGAGT | Chatzinikolaou et al., 2017 |
| | Rev: | TAGGGAGAGCGGCTTTTACA | |
| *Col1a2* | Fwd: | AAAGAGAAGGATTGGTCAGAGCAGT | Varrault et al., 2017 |
| | Rev: | GCCAAGGGAGGAGACTTAGTTG | |
| *Col9a2* | Fwd: | CTCTGGACTTATTTTTATTGGTATCTTTT | Varrault et al., 2017 |
| | Rev: | CAGGGAAGATGGATGTTTAAATACTG | |

***Source of published primers**:*


Chatzinikolaou, G. et al. (2017) ERCC1-XPF cooperates with CTCF and cohesin to facilitate the developmental silencing of imprinted genes. **Nat Cell Biol** 19, 421-432.

Leeb, M. et al. (2010) Polycomb complexes act redundantly to repress genomic repeats and genes. **Genes Dev** 24, 265-276.

Sanli, I. et al. (2018) Meg3 non-coding RNA expression controls imprinting by preventing transcriptional upregulation in cis. **Cell Rep** 23, 337-348.

Varrault, A. et al. (2017) Mouse parthenogenetic embryonic stem cells with biparental-like expression of imprinted genes generate cortical-like neurons that integrate into the injured adult cerebral cortex. **Stem Cells** 36, 192-205.

### ***Fosmid and BAC probes for FISH experiments***

All coordinates GRCm38/mm10

| **Fosmid** | **Locus** | **Position** | **Length (bp)** |
|---|---|---|---|
| WIBR1-1062J20 | mHIDAD CTCF | chr7:142,316,692-142,353,854 | 37,162 |
| WIBR1-090J20 | *H19* | chr7:142,579,083-142,621,420 | 42,337 |
| WIBR1-399J17 | *Igf2* | chr7:142,656,682-142,695,317 | 38,635 |
| WIBR1-1319N18 | Upstream *Dlk1* | chr12:109,363,854-109,403,919 | 40,065 |
| WIBR1-1116K16 | *Dlk1* | chr12:109,435,989-109,471,592 | 35,603 |
| WIBR1-1703L18 | *Meg3* | chr12:109,539,156-109,578,782 | 39,626 |
| WIBR1-2409H13 | *Dio3* | chr12:110,254,267-110,294,032 | 39,765 |

| **BAC** | **Locus** | **Position** | **Length (bp)** |
|---|---|---|---|
| RP23-119N23 | Upstream *Begain* | chr12:108,816,610-109,016,762 | 200,153 |
| RP23-421C16 | *Meg3 / Gtl2* | chr12:109,480,824-109,658,133 | 177,310 |
| RP23-41I14 | Downstream *Dio3* | chr12:110,431,516-110,659,759 | 228,244 |
